# STRIP2 promotes colorectal cancer malignant progression via stabilizing LCN2 to suppress ferroptosis

**DOI:** 10.64898/2026.05.16.725308

**Authors:** Chunxiao Hu, Xiaohua Ye, Xi Chen, Shengyue Zhou, Honghua Hu, Jin Ding, Weijun Teng

## Abstract

Colorectal cancer (CRC) presents a significant global health challenge with high incidence rates, mortality, and poor five-year survival rates, particularly in advanced cases, despite the availability of existing treatments. This study investigates the role of Striatin-interacting protein 2 (STRIP2) in CRC progression and the underlying mechanisms. To evaluate the impact of STRIP2 on CRC, we conducted *in vitro* experiments to assess cell proliferation, migration, invasion, and apoptosis in CRC cell lines. Furthermore, *in vivo* studies were performed utilizing nude mouse models. Transcriptomic analysis, Western blotting, co-immunoprecipitation assays, and ferroptosis validation were employed to elucidate the role of STRIP2 in CRC. The findings indicate that STRIP2 is overexpressed in CRC and drives malignant behaviors in both *in vitro* and *in vivo* settings. Overexpressing STRIP2 (oe-STRIP2) suppresses ferroptosis, leading to increased cell proliferation and reduced oxidative stress, whereas silencing STRIP2 (si-STRIP2) has the opposite effect. Mechanistically, STRIP2 stabilizes LCN2, an effector downstream of IL-17, by preventing its K48-linked ubiquitination and subsequent degradation. This stabilization enhances the anti-ferroptosis capacity of CRC cells, thereby promoting cell survival and facilitating malignant progression in CRC. These findings suggest STRIP2 as a promising diagnostic biomarker and therapeutic target for CRC intervention and warrant further investigation.

**Highlight:**

- STRIP2 is overexpressed in CRC tissues and cells
- STRIP2 drives CRC malignant phenotypes both *in vitro* & *in vivo*
- STRIP2 blocks LCN2 Ubiquitination and stabilizes LCN2
- STRIP2 promotes CRC Malignancy via LCN2 stabilization and ferroptosis suppression
- STRIP2 could be a potential biomarker and therapeutic target for CRC

## Introduction

Colorectal cancer (CRC) remains a significant global health challenge, being the third most common cancer, accounting for approximately 10% of all new cancer cases worldwide, and the second leading cause of cancer-related deaths worldwide (1, 2). For instance, global new CRC cases are projected to rise from 1.9 million in 2020 to 3.2 million in 2040 (3, 4). Despite current treatments for CRC, including surgery, chemotherapy, radiotherapy, and targeted therapy, the 5-year overall survival rate for advanced CRC is only 10% to 15% (5, 6).

Adenocarcinoma, the most common CRC subtype, typically has an indolent course, with a 5–10-year window between the development of early adenomas and malignant transformation (7). The multistep malignant transformation of colorectal epithelial cells is driven by successive oncogenic activation, tumor suppressor inactivation, and dysregulation of post-translational modifications (7); exploring novel driver genes and their downstream regulatory cascades is essential for uncovering effective anti-CRC therapeutic targets.

Striatin-interacting protein 2 (STRIP2, FAM40B), a core constituent of the STRIPAK complex, participates in cytoskeleton remodeling (8) and multiple kinase/phosphatases signalling regulation (9, 10). Cumulative evidence has verified aberrant upregulation of STRIP2 in lung (11, 12), gastric cancer (13), and other digestive malignancies. STRIP2 is also involved in embryonic stem cell differentiation and atherosclerosis (14, 15). Notably, CRC patients with elevated STRIP2 expression exhibit reduced overall, disease-specific, and progression-free survival (16). Despite this association, the molecular mechanisms by which STRIP2 drives malignant progression in CRC remain unclear. In this study, we reveal that STRIP2 stabilizes the IL-17 downstream effector Lipocalin 2 (LCN2), thereby inhibiting ferroptosis and promoting CRC malignant phenotypes *in vitro* and *in vivo*.

Ferroptosis is an iron-dependent form of cell death characterized by overload of intracellular labile iron, depletion of glutathione (GSH), and uncontrolled lipid peroxidation (17). Ferroptosis is executed through three main processes: first, excess free iron initiates the Fenton reaction, generating reactive oxygen species (ROS) that trigger phospholipid peroxidation (18). Second, the System Xc, solute carrier family 7-member 11 (SLC7A11), GSH, and glutathione peroxidase 4 (GPX4) antioxidant axis serves as the primary line of defense against the accumulation of lipid peroxides (19). Third, the esterification of polyunsaturated fatty acids mediated by acyl-CoA synthetase long-chain family member 4 (ACSL4) provides substrates for peroxidation, thereby sensitizing cells to ferroptosis (20). Additionally, FSP1 and NRF2 represent alternative anti-ferroptotic pathways that operate independently of GPX4 (21). In CRC, the suppression of endogenous ferroptosis is a well-recognized characteristic of malignancy (22). This allows tumor cells to evade oxidative damage, develop resistance to chemotherapy, and maintain metastatic colonization (23).

LCN2 is an iron-binding protein that inhibits ferroptosis in CRC (24). It stabilizes intracellular iron to prevent the Fenton reaction, while also upregulating antioxidant genes such as SLC7A11, GPX4, FSP1, and NRF2 and downregulating ACSL4 to block lipid peroxidation (25). Our transcriptomic screening revealed significant downregulation of LCN2 after STRIP2 depletion. This study shows that STRIP2 inhibits K48-linked ubiquitination and proteasomal degradation of LCN2, thus preventing ferroptosis and promoting CRC progression *in vitro* and *in vivo*. STRIP2 has been implicated in gastrointestinal cancer progression, but its role in modulating ferroptosis via LCN2 has never been explored. This study fills this gap by identifying STRIP2 as a novel post-translational stabilizer of LCN2 that drives ferroptosis resistance and CRC malignancy, and by identifying a potential biomarker and therapeutic target for CRC treatment.

## Materials and methods

### Bioinformatics analysis of web databases

All public dataset analyses were performed using R software (v4.2.1; R Foundation for Statistical Computing, Vienna, Austria; https://www.r-project.org/) following standard package documentation, with all database access completed on 12th Dec. 2025. All human genomic data were mapped to the GRCh38.p13 human reference genome assembly.

First, the GSE44861 microarray dataset was retrieved from the Gene Expression Omnibus (GEO, https://www.ncbi.nlm.nih.gov/geo/), containing transcriptomic profiles of 56 primary CRC tissues and 55 paired adjacent non-colon mucosal tissues from human CRC patients. Raw expression matrices were imported into the DESeq2 R package (v1.36.0) for normalisation and differential expression gene (DEG) identification. DEG screening thresholds were defined as adjusted P (Padj) < 0.05 and absolute log fold change (|log FC|) ≥ 1. This analysis yielded 241 DEGs, comprising 74 upregulated and 167 downregulated transcripts; STRIP2 was significantly overexpressed in colon and rectal cohorts relative to matched non-mucosa.

Secondly, pan-cancer and CRC cell line STRIP2 expression data were extracted from the Human Protein Atlas (HPA, https://www.proteinatlas.org/), a public human tissue and cell line protein expression database, to compare endogenous STRIP2 abundance across all major human malignancies and standard CRC in vitro models.

Thirdly, RNA-seq raw counts for colon adenocarcinoma (COAD) and rectal adenocarcinoma (READ) cohorts were downloaded from The Cancer Genome Atlas (TCGA, https://portal.gdc.cancer.gov/). The Linear Models for Microarray and Omics Data (limma) R package (v3.52.4) was utilised for normalisation and intergroup comparison of STRIP2 mRNA levels between histologically confirmed specimens and non-malignant colorectal mucosal controls.

### Patients and clinical specimens

Five pairs of primary human CRC tissues and matched adjacent non-colorectal mucosal tissues were prospectively collected from patients undergoing curative primary CRC resection at the Affiliated Jinhua Hospital, Zhejiang

University School of Medicine in 2024. Exclusion criteria included neoadjuvant chemotherapy, radiotherapy, or immunotherapy administered prior to surgical resection. All tissue diagnoses were independently validated by two board-certified gastrointestinal pathologists via routine haematoxylin and eosin (H&E) staining. Adjacent non-mucosa was harvested ≥5 cm distal from the macroscopic margin and confirmed histologically free of dysplasia or malignant infiltration.

Each tissue specimen was bisected immediately post-resection: one tissue fragment was snap-frozen in liquid nitrogen and stored long-term at −80°C for protein extraction and Western blotting; the second fragment was fixed in 4% paraformaldehyde (PFA; Solarbio, Beijing, China, Cat# P1110) for 24 h at room temperature, sequentially dehydrated in graded ethanol, cleared in xylene, and embedded in paraffin wax for immunohistochemistry (IHC) staining. Patient clinical and pathological data (age, gender, differentiation, and TNM stage) are summarised in Table S1.

This human tissue study received formal ethical approval from the Committees for Ethical Review at the Affiliated Jinhua Hospital, Zhejiang University School of Medicine (Approval Reference No. 20252220101). Written informed consent was obtained from all enrolled participants prior to tissue acquisition in compliance with the Declaration of Helsinki.

### Cell lines and cell culture

Human CRC cell lines HCT116 (Cat# TCHu 99), SW480 (Cat# TCHu172) and human embryonic kidney (HEK) 293T/17 (Cat# GNHu44) were obtained from the Chinese Academy of Sciences cell bank (Shanghai, China). All cell lines were confirmed mycoplasma-negative via the MycoBlue Mycoplasma Detection Kit (Vazyme, Nanjing, China, Cat# D101-01) prior to all functional assays.

Cells were cultured in the cell line-specific media: SW480 cells in Leibovitz’s L-15 medium (Gibco, Thermo Fisher Scientific, Waltham, MA, USA, Cat# 11415064); HCT116 cells in McCoy’s 5A medium (Gibco, Cat# 16600082); and 293T/17 cells in Dulbecco’s Modified Eagle Medium (DMEM; Gibco, Cat# 11965092).

All base media were supplemented with 10% foetal bovine serum (FBS; Vazyme, Cat# F101), 100 IU/ml penicillin G sodium and 100 µg/ml streptomycin sulphate (Solarbio, Cat# P1400). Cells were cultured in a humidified incubator (Thermo Scientific Heracell 150i, Thermo Fisher Scientific) maintained at 37°C with 5% CO. HCT116 was selected as the primary in vitro model for xenograft animal studies, RNA-seq profiling, LCN2 protein stability assays, and ferroptosis functional validation due to due to its model stability, experimental sensitivity, and clinical relevance.

### RNA extraction and quantitative real-time PCR

Total RNA was extracted using TRIquick Total RNA Extraction Reagent (Solarbio, Cat# R1100). RNA purity and concentration were quantified via NanoDrop One UV-Vis spectrophotometer (Thermo Fisher Scientific). First-strand complementary DNA (cDNA) was synthesised from 1 µg total RNA per sample using the PrimeScript™ RT Reagent Kit (Takara, Tokyo, Japan, Cat# RR037A).

Quantitative real-time PCR (qPCR) was performed using the SYBR qPCR Master Mix Kit (Vazyme, Cat# Q311) on a fluorescence quantitative real-time PCR detection system (CFX Opus, Bio-Rad, Hercules, CA, USA). Primer sequences for target genes and the endogenous reference glyceraldehyde-3-phosphate dehydrogenase (*GAPDH*) are provided in Table S2. Each 20 µl reaction system contained 10 µl 2× SYBR Master Mix, 10 µM forward primer and 10 µM reverse primer, 2 µl diluted cDNA template, and nuclease-free water. Thermal cycling parameters: initial denaturation at 95°C for 3 min; 40 amplification cycles of 95°C for 10 s, 60°C annealing/extension for 30 s; melting curve analysis (95°C 15 s, 60°C 60 s, ramp to 95°C) to validate amplicon specificity. All samples were assayed in three technical replicates per biological replicate. Relative gene expression was calculated using the comparative 2^−ΔΔCq method, with *GAPDH* serving as the constitutively expressed internal reference gene.

### STRIP2 overexpression and knockdown shRNA vectors construction and cell transfection

Full-length human STRIP2 coding sequence (CDS) was amplified from HCT116 cDNA using Phanta Super-Fidelity DNA Polymerase (Vazyme, Cat# P501-d3). Amplicons were digested with restriction enzymes and ligated into two independent expression vectors: pHAGE-CMV-MCS-IRES-ZsGreen lentiviral vector (Novopro, Shanghai, China, Cat# V014630) and pcDNA3.1 eukaryotic expression vector (Thermo Fisher Scientific, Waltham, MA, USA). Empty pcDNA3.1 vector was transfected as the overexpression negative control (oe-NC), and the recombinant pcDNA3.1-STRIP2 construct was designated oe-STRIP2.

STRIP2-targeted short hairpin RNA (shRNA) was designed via Invitrogen Block-iT RNAi Designer online tool (<u>Invitrogen Block-iT RNAi Designer</u>). The sense strand sequence was 5′-AGACCTTCCGCACTGAATTAA-3′. Oligonucleotides were synthesised by GeneralBio (Anhui, China), annealed, and subcloned into pcDNA3.1 to generate pcDNA3.1-shSTRIP2 (si-STRIP2). A non-targeting scrambled shRNA sequence inserted into pcDNA3.1 served as the knockdown negative control (shNC/si-NC). For lentiviral stable cell line generation, the shSTRIP2 cassette was packaged into lentiviral particles with psPAX2 (Addgene, Watertown, MA, USA, Cat.# 12260) and pMD2.G (Addgene, Cat.# 12259) packaging plasmids in HCT116 cells or HEK 293T/17 cells.

All transient transfections were performed in 6-well culture plates when cells reached 70–80% confluence using Lipofectamine 3000 Transfection Reagent (Invitrogen, Thermo Fisher Scientific, Cat# L3000015). Plasmid-lipid complexes were prepared in Opti-MEM reduced serum medium (Invitrogen, Cat# 31985070), mixed gently, and incubated at room temperature for 30 min to enable complex formation. Complex suspensions were added dropwise to adherent cells; culture medium was refreshed 6 h post-transfection. Cells were harvested 48 h post-transfection for all downstream molecular and functional assays.

### Cell proliferation assays

Cell proliferation was evaluated using CCK-8 assay, EdU incorporation assay and colony formation assay, each assay with six biological replicates per group. Data are presented as mean ± standard deviation (SD).

### CCK-8 assay

CCK-8 cell counting kit (Vazyme, Cat# A311-01) was applied per manufacturer instructions. Absorbance at 450 nm was measured via a SpectraMax iD3 microplate reader (Molecular Devices, San Jose, CA, USA) at predefined time points to quantify viable cell density.

### EdU incorporation assay

5-Ethynyl-2’-deoxyuridine (EdU) labelling was conducted using VaClick 488-EdU Cell Proliferation Test Kit (Vazyme, Cat# A411-01 for Figure 2D) and Cell-Light EdU Apollo567 In Vitro Kit (RiboBio, Guangzhou, China, Cat #C10310-1 for Figure 7B, 7E). EdU-positive proliferating cells were imaged under an inverted fluorescence microscope (Shanghai Cewei Optoelectronic Technology, model LWD300-38LFT) and quantified using ImageJ software v1.53j. The EdU incorporation rate was calculated as the ratio of the number of EdU-incorporated cells to the number of Hoechst 33342-staining cells.

### Colony formation assay

A total of 5,000 HCT116 or SW480 cells were seeded per 6-well plate and cultured for 14 days with complete medium refreshed every 72 h. Colonies were fixed in 4% PFA, stained with 0.5% crystal violet (Solarbio, Cat# G1063) for 30 min at room temperature, and washed three times with phosphate-buffered saline (PBS; pH 7.4). Colonies containing ≥50 cells were counted under brightfield microscopy.

### Cell migration assay

The wound-healing assay was conducted to evaluate cell migration. Cells were seeded in 6-well plates and cultured to 90-100% confluent monolayers. A uniform linear scratch wound was generated across each well using a sterile 200 μl pipette tip; detached floating cells were washed twice with PBS. Culture medium was replaced with serum-free base medium to eliminate proliferation-driven wound closure. Brightfield images at 10× objective magnification were captured at 0 h, 24 h and 48 h post-scratch using an inverted microscope (ICX41, Sunny Optical, Zhejiang, China). Wound closure area was quantified in ImageJ v1.53j. Three independent biological replicates were performed for each group.

### Cell invasion assay

Matrigel-precoated transwell cell culture inserts (8 μm pore size; Corning, Corning, NY, USA) were used to assess invasive capacity. A total of 1 × 10 si-NC or si-STRIP2 HCT116/SW480 cells were suspended in serum-free medium and seeded into the upper transwell chamber; the lower chamber contained complete medium supplemented with 10% FBS as a chemotactic attractant. After 24 h incubation at 37°C, non-invasive cells remaining on the upper membrane surface were mechanically removed with cotton swabs. Invaded cells on the lower membrane were fixed, stained with 0.5% crystal violet, and imaged under 10× magnification. Total invasive cell counts were quantified via ImageJ v1.53j across three biological replicates. Parallel whole-cell lysates were collected for Western blot analysis of epithelial-mesenchymal transition (EMT) marker proteins (E-cadherin, vimentin and snail) expression.

### Cell apoptosis assay

Apoptosis was quantified via Annexin V-FITC/propidium iodide (PI) dual staining using the Annexin V-FITC/PI Apoptosis Detection Kit (Vazyme, Cat# A211-01). Harvested trypsinised cells were washed with pre-chilled PBS, resuspended in 1× binding buffer, incubated with Annexin V-FITC and PI staining reagents for 15 min at room temperature in the dark, and analysed immediately on a Novocyte Advanteon Flow Cytometer System (Agilent Technologies, Santa Clara, CA, USA). Raw cytometry data were processed with FlowJo v10 software (FlowJo LLC, Ashland, OR, USA). The proportion of early apoptotic (Annexin V PI), late apoptotic (Annexin V PI) and necrotic (Annexin V PI) populations within gated intact single cells was quantified across three biological replicates. Complementary Western blot analysis of apoptosis effector proteins (BCL2, BAX, Pro-Caspase-3, cleaved Caspase-3) was performed to validate flow cytometry results.

### In-House RNA-seq transcriptomics analysis

Total RNA extracted from stably transducted si-NC and si-STRIP2 HCT116 cells (n = 3) was submitted to GeneralBio Co., Ltd. (Anhui, China) for mRNA sequencing on the Illumina NovaSeq 6000 platform (Illumina, San Diego, CA, USA). Briefly, 1 µg total RNA per sample was enriched for poly(A) mRNA using oligo(dT) magnetic beads; strand-specific mRNA libraries were constructed with the NEBNext Ultra II Directional RNA Library Prep Kit (New England Biolabs, Ipswich, MA, USA, Cat# E7760S) following the manufacturer’s instruction. Raw FASTQ sequencing reads were quality-filtered to generate clean data (Clean Data) and aligned to the GRCh38.p13 human reference genome using HISAT2 (v2.2.1) with default alignment parameters. Gene-level read counts were quantified via HTSeq (v0.6.1) and normalised to Fragments Per Kilobase of transcript per Million mapped reads (FPKM). DEG calling was performed with DESeq2 (v1.36.0) using identical thresholds (Padj < 0.05, |log FC| ≥ 1).

Functional enrichment analyses were executed via clusterProfiler (v4.4.4) and org.Hs.eg.db (v3.15.0) R packages to annotate DEGs against Kyoto Encyclopedia of Genes and Genomes (KEGG) pathway databases. Gene Set Enrichment Analysis (GSEA) was run using Hallmark curated gene sets from the Molecular Signatures Database (MSigDB v7.5.1) to identify perturbed oncogenic and metabolic signalling axes. A gene expression heatmap visualising top variable DEGs was plotted with the pheatmap (v1.0.12) R package.

### In vivo nude mouse xenograft and liver metastasis models

Male BALB/c athymic nude mice (4–5 weeks old, body weight 18–20 g) were purchased from Ziyuan Laboratory Animal Technology Co., Ltd. (Hangzhou, China). All animals were housed in a specific-pathogen-free facility with controlled environmental conditions: ambient temperature 22 ± 2 °C, relative humidity 50–60%, 12 h light/dark diurnal cycle. All in vivo animal experimental protocols were approved by the Experimental Animal Ethics Committee of Jinhua Institute of Food and Drug Inspection and Testing (Approval Reference No. AL-JSYJ202473). All animal handling, anaesthesia, surgery and euthanasia procedures strictly complied with the ARRIVE 2.0 reporting guidelines for animal research.

A preliminary pilot study (n = 4 mice per group) was conducted to establish growth and metastatic burden baseline values for power analysis. Two independent *in vivo* models (subcutaneous tumorigenesis and splenic injection liver metastasis) were each performed across three consecutive identical experimental cohorts (n = 12 mice per cohort; n = 6 per group with stably expressing control shRNA (sh-NC) or STRIP2-targeted shRNA (sh-STRIP2) HCT116 cells). Investigators performing measurement, tissue harvesting and histological quantification were fully blinded to group allocation. The study was powered at α = 0.05, statistical power (1−β) = 0.8 to detect a 30% relative difference in primary volume and hepatic metastatic burden between groups.

### Subcutaneous xenograft assay

Each cohort comprises 12 mice, randomized by body weight into two groups (sh-NC and sh-STRIP2 HCT116 cells; n = 6 per group). A single-cell suspension (5 × 10 cells in 200 μl PBS) was injected subcutaneously into the right dorsal flank. Tumor dimensions were measured with digital calipers every 3 days, and volume was calculated as V = 0.5 × length × width². On day 27, mice were deeply anesthetized under 2% isoflurane deep anesthesia (RWD Life Science, Shenzhen, China). Whole blood was collected via cardiac puncture into anticoagulant-free 1.5 ml tubes, clotted at room temperature and centrifuged. Serum was harvested, aliquoted, stored at −80°C, and used for enzyme-linked immunosorbent assay (ELISA). Mice were euthanized by cervical dislocation after blood collection. Tumors were excised, weighed, photographed, and fixed in 4% paraformaldehyde for 24 h and paraffin-embedded for hematoxylin and eosin (H&E) and Ki-67 IHC.

### Splenic injection liver-metastasis model

Mice were anaesthetised via 2% isoflurane inhalation, and a 0.5 cm subcostal laparotomy incision was made to expose the spleen. A 29-gauge insulin syringe was used to inject 5 × 10 HCT116 shNC or shSTRIP2 cells suspended in 200 μl sterile PBS into the splenic lower pole. Gentle digital compression was applied for 1 min post-injection to achieve haemostasis before layered abdominal wall and skin closure. Five weeks after cell administration, all animals were euthanised, and whole livers were harvested and imaged under a stereomicroscope to count macroscopic surface metastatic nodules. The largest hepatic lobe was coronally sectioned, FFPE-processed, and serial 4 μm histological slices cut at 200 μm intervals for H&E staining. Total metastatic lesion area was quantified from digital slide scans using ImageJ v1.53j by two independent blinded pathologists.

### Enzyme-linked immunosorbent assay

Mice serum carcinoembryonic antigen (CEA) and carbohydrate antigen 19-9 (CA199) levels were quantified using ELISA kits (Coibo Biotechnology, Shanghai, China): Mouse CEA ELISA Kit (Cat# CB13612-Mu) and Mouse CA199 ELISA Kit (Cat# CB10990-Mu) according to manufacturer’s instructions. Absorbance readings at 450 nm were acquired on a SpectraMax iD3 microplate reader (Molecular Devices) for standard curve interpolation of antigen concentrations.

### Protein extraction and Western blotting

Adherent cells were washed twice with ice-cold PBS and lysed for 30 min on ice in complete RIPA buffer (Beyotime, Shanghai, China, Cat# P0013B) supplemented with 1 mM phenylmethylsulfonyl fluoride (PMSF; Solarbio, Cat# P0100) and 1× protease/phosphatase inhibitor cocktail (Beyotime, Cat# P1045). Cell lysates were clarified by centrifugation at 12,000 × g for 15 min at 4°C. Total soluble protein concentration was determined via the bicinchoninic acid (BCA) colorimetric assay (Beyotime, Cat# P0010), with absorbance measured at 562 nm on a microplate reader. Thirty micrograms of total protein per sample were mixed with 5× protein loading buffer, denatured at 95°C for 10 min, and separated by 10% sodium dodecyl sulphate-polyacrylamide gel electrophoresis (SDS-PAGE) at constant voltage. Separated proteins were wet-electrotransferred onto 0.22 μm polyvinylidene difluoride (PVDF) membranes (Millipore, Burlington, MA, USA, Cat# ISEQ00010). Membranes were blocked with 5% (w/v) non-fat dried milk diluted in Tris-buffered saline containing 0.1% Tween-20 (TBST) for 1 h at room temperature to saturate non-specific antibody binding sites. Blocked membranes were incubated overnight at 4°C with primary antibodies (all from Proteintech, Wuhan, China) diluted in TBST. Antibody name, catalogue number and dilution factor are listed in Table S3. Membranes were washed three times for 10 min each in TBST, then incubated with horseradish-peroxidase (HRP)-conjugated secondary antibodies for 1 h at room temperature: goat anti-rabbit IgG (FUDE BIOLOGICA, Hangzhou, China, Cat# FDK9001, 1:10,000 dilution) and goat anti-mouse IgG (FUDE BIOLOGICA, Cat# FDK9002, 1:10,000 dilution). Immunoreactive protein bands were visualised using Amersham ECL Prime Western Blotting Detection Reagent (Cytiva, Chicago, IL, USA, Cat# RPN2232). Chemiluminescent imaging was performed using an SCG-W2000 system (Wuhan Sevier Biotechnology Co., Ltd., Wuhan, China) and quantified using ImageJ v1.53j; GAPDH served as the loading control.

### Immunohistochemical staining

Paraffin sections underwent antigen retrieval in 0.01 M citrate buffer (pH 6.0) at 98°C for 20 min. Endogenous peroxidase was quenched with 3 % hydrogen peroxide (HO) for 10 min. Non-specific antibody binding was blocked with 5% normal goat serum (Solarbio, Cat# SL038) for 30 min. Slides were incubated overnight at 4°C with primary anti-STRIP2 or anti-Ki-67 antibody (Proteintech, Table S3). After three PBS washes, HRP-conjugated goat anti-rabbit polymer secondary reagent (Gene Tech, Shanghai, China, Cat# GK500710) was applied for 30 min at room temperature. Chromogenic signals were developed using 3,3’-diaminobenzidine (DAB) substrate, and slides were counterstained with haematoxylin for nuclear visualisation. Negative control slides were processed identically with primary antibody omitted to confirm staining specificity.

Slides were digitally scanned using an Aperio AT2 whole-slide scanner (Leica Biosystems). Two independent blinded pathologists quantified STRIP2 and Ki-67 immunoreactivity using the H-score system (range 0–300; calculated as percentage positive cells × staining intensity). H-scores ≥50 were defined as high STRIP2/Ki-67 expression.

### Protein stability assays

MG132 proteasome inhibition assay and cycloheximide (CHX) chase assay were used to evaluate protein stability.

### MG132 proteasome inhibition assay

Stable sh-STRIP2 293T/17 cell lines were generated via lentiviral transduction followed by 72 h puromycin selection (2 μg/ml; Solarbio, Cat# P8230) and Western blot validation of STRIP2 knockdown efficiency. Cells were seeded at 2 × 10 cells per 6-well and cultured to 70–80% confluence before being divided into four treatment groups: vehicle control, oe-STRIP2 only, MG132 only, and oe-STRIP2 + MG132. The vehicle control group was cultured with complete medium containing 0.1% dimethyl sulfoxide (DMSO) for 24 h. The oe-STRIP2-only group was transfected with pcDNA3.1-STRIP2 and cultured for 24 h post-transfection without additional drugs. The MG132-only group was exposed to 10 μM MG132 (MedChemExpress (MCE), Shanghai, China, Cat# HY-13259) for 4 h. The oe-STRIP2 + MG132 group was transfected with pcDNA3.1-STRIP2, cultured for 20 h post-transfection, then co-incubated with 10 μM MG132 for 4 h (total 24 h). Total cellular proteins were extracted and processed for subsequent Western blot to detect target protein expression.

### CHX chase assay

si-NC and si-STRIP2 HCT116 cells were treated with 100 μg/ml CHX (MCE, Cat# HY-12320) to block de novo protein synthesis. Total cellular protein was extracted at 0, 3, 6, 9, 12, and 15 h post-CHX treatment. Western blot was used to quantify temporal LCN2 protein degradation kinetics in STRIP2-knockdown vs control cells.

### Co-immunoprecipitation Assay

HCT116 cells were co-transfected with Flag-tagged STRIP2, HA-tagged LCN2, and HA-tagged ubiquitin (HA-Ub) expression plasmids. At 48 h post-transfection, cells were lysed on ice in complete RIPA lysis buffer supplemented with 1× protease inhibitor cocktail (Beyotime, Cat# P1005). Clarified whole-cell lysates were retained as input controls; remaining lysate was incubated with anti-HA antibody-conjugated agarose beads at 4°C with gentle rotation overnight to capture HA-labelled protein complexes. Beads were washed five times with ice-cold RIPA buffer to eliminate non-specific binding. Immunoprecipitated protein complexes were eluted, separated by SDS-PAGE, and immunoblotted with primary antibodies targeting HA, LCN2, and STRIP2; GAPDH was used as an input loading control.

### Ferroptosis validation

To validate the link between STRIP2 and ferroptosis regulation in CRC. oe-STRIP2 cells were treated with 30 μM erastin ferroptosis inducer (MCE, Cat# HY-15763) or 30 μM erastin and 2 μM ferrostatin-1 (Fer-1) ferroptosis inhibitor (MCE, Cat# HY-100579) for 24 h; while si-STRIP2 cells were treated with 2 μM Fer-1 for 24 h; with vehicle DMSO treatment serving as parallel controls. The following ferroptosis-related assays were measured across three biological replicates: (1) Western blot of ferroptosis regulatory proteins: pro-ferroptotic ACSL4; anti-ferroptotic SLC7A11, GPX4, FSP1, and NRF2. (2). CCK-8 and EdU proliferation assays were used to measure ferroptosis-mediated growth suppression. (3). Intracellular glutathione (GSH, key indicator of cellular antioxidant capacity) and malondialdehyde (MDA, lipid peroxidation marker linked to ferroptosis) were measured using commercial colorimetric kits (Geruisi Bio, Suzhou, China; GSH assay kit Cat# G0206W, MDA assay kit Cat# G0109W). (4). Lipid ROS in tumor cells was detected using a lipid peroxidation assay kit containing BODIPY 581/591 C11 (Beyotime, Shanghai, China, Cat. #S0043S). Treated tumor cells were incubated with 2 μM C11 BODIPY in the dark at 37°C under 5% CO for 30 min. After three washes with ice cold PBS, green/red fluorescence signals of C11 BODIPY (Ex 488 nm / Em 510 nm and Ex 581 nm / Em 590 nm) were acquired using a fluorescence microscope. (5). Intracellular labile iron levels were measured using FerroOrange (Dojindo, Kumamoto, Japan, Cat. #F374). HCT116 cells were seeded in culture dishes and incubated for 24 h. Adherent cells were rinsed twice with pre-warmed serum-free medium, then stained with 0.5 μM FerroOrange for 30 min at 37°C in the dark. After washing twice with pre-cooled PBS, orange fluorescence intensities were measured at Ex 543 nm / Em 580 nm with a multifunctional microplate reader.

### Statistical analysis

Statistical analyses were conducted using GraphPad Prism v10.0 software (GraphPad Software, San Diego, CA, USA): unpaired two-tailed Student’s independent samples t-test was used for comparisons between two groups, and one-way ANOVA with Tukey’s post-hoc test was used for multiple groups. Statistical significance was defined as P < 0.05 (*P < 0.05, **P < 0.01, ***P < 0.001).

## Results

### STRIP2 is overexpressed in colorectal cancer tissues and cell lines

To investigate the clinical significance of STRIP2 in CRC, we first profiled its expression across pan-cancer datasets. STRIP2 was found to be broadly upregulated in multiple types of human malignancies (Fig. 1A). In addition, STRIP2 was ubiquitously expressed across a panel of 63 CRC cell lines with varying differentiation statuses (Fig. 1B). RNA-Seq analysis using the limma R package on the TCGA_COAD and TCGA_READ datasets reinforced that STRIP2 is significantly overexpressed in both colon and rectal cancers (Fig. 1C). Western blot analyses (Fig. 1D) showed that STRIP2 was upregulated in CRC tissues compared to paired non-tumor tissues. IHC analysis (Fig. 1E) demonstrated that STRIP2 protein expression was elevated in tumor tissues compared to paired non-tumor tissues, with distinct cytoplasmic and nuclear localization. These findings indicate that STRIP2 is frequently upregulated in CRC and may represent a potential diagnostic biomarker.

**Figure 1.**
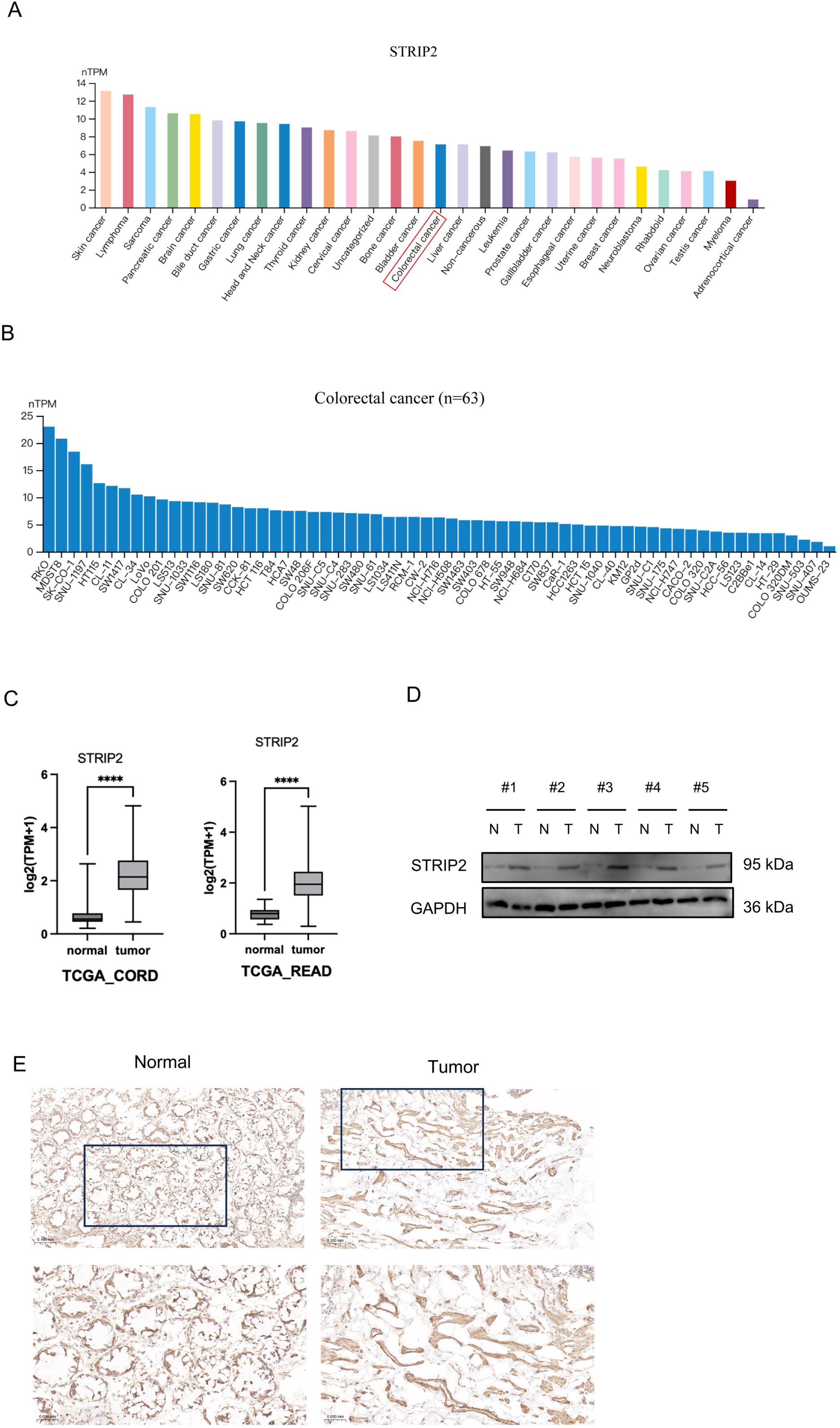
STRIP2 is overexpressed in CRC tissues and cell lines. (A) Pan-cancer analysis of STRIP2 mRNA expression across multiple human cancer types. (B) Relative STRIP2 mRNA levels in a panel of CRC cell lines. (C) Relative STRIP2 mRNA expression in normal colon/rectal tissues and CRC samples from the TCGA-COAD and TCGA-READ datasets. (D) Western blot analysis of STRIP2 protein levels in paired CRC tumor (T) and matched adjacent normal (N) tissues (n=5). (E) Immunohistochemical (IHC) staining of STRIP2 protein in paired CRC tumor (T) and matched adjacent normal (N) tissues (n=5). Scale bars: 50 μm (left), 100 μm (right). Data are presented as mean ± SD. ***P<0.001. STRIP2: striatin-interacting phosphatase and kinase subunit 2; CRC: colorectal cancer; TCGA: The Cancer Genome Atlas; COAD: colon adenocarcinoma; READ: rectum adenocarcinoma; IHC: immunohistochemistry.

### STRIP2 knockdown suppresses CRC cell proliferation

To explore the biological function of STRIP2 in CRC, we silenced STRIP2 expression in SW480 and HCT116 cells using small interfering RNA (siRNA). Both qRT-PCR and Western blot analyses verified robust knockdown efficiency at the mRNA and protein levels in both cell lines (Fig. 2A, B).

**Figure 2.**
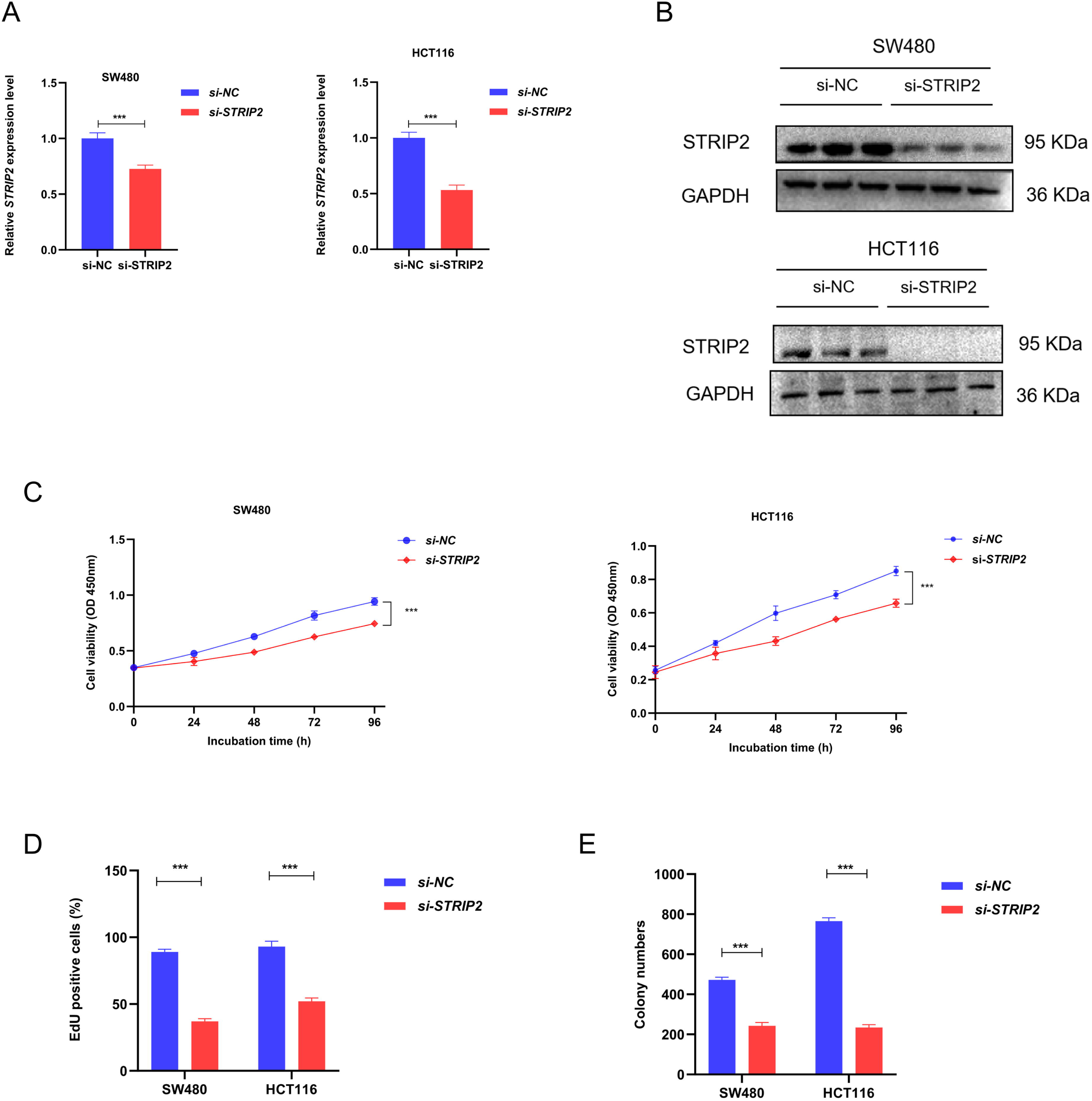
STRIP2 silencing inhibits CRC cell proliferation. (A) qRT-PCR validation of STRIP2 knockdown efficiency in SW480 and HCT116 cells (n=3). (B) Western blot analysis confirming STRIP2 protein silencing in SW480 and HCT116 cells (n=3). (C–E) Proliferation functional assays in SW480 and HCT116 cells (n=6): (C) CCK-8 cell viability assay, (D) EdU incorporation assay, (E) Colony formation assay. Data are presented as mean ± SD. ***P<0.001. STRIP2: striatin-interacting phosphatase and kinase subunit 2; CRC: colorectal cancer; si: silencing; siRNA: small interfering RNA; NC: negative control; qRT-PCR: quantitative real-time PCR; CCK-8: Cell Counting Kit-8; EdU: 5-ethynyl-2′-deoxyuridine.

We then assessed the impact of STRIP2 depletion on CRC cell proliferation. CCK-8 assays demonstrated that STRIP2 knockdown significantly reduced cell viability in a time-dependent manner in both SW480 and HCT116 cells (all P < 0.001, Fig. 2C). EdU incorporation assays consistently revealed a marked reduction in the proportion of actively proliferating cells following STRIP2 silencing (Fig. 2D). Colony formation assays showed the number of colonies were dramatically decreased in the si-STRIP2 groups compared with the si-NC control groups (Fig. 2E). These findings collectively demonstrate that STRIP2 is required for sustaining the proliferative capacity of CRC cells.

### STRIP2 promotes migration and invasion and inhibits apoptosis in CRC cells

We next evaluated the role of STRIP2 in CRC cell motility and invasiveness. Wound-healing assays showed that STRIP2 knockdown significantly delayed wound closure at 24 h and 48 h in both SW480 and HCT116 cells, indicating impaired migratory ability (Fig. 3A, B). Transwell invasion assays further showed that STRIP2 depletion drastically reduced the number of cells that penetrated Matrigel-coated membranes, reflecting suppressed invasive potential (all P < 0.001, Fig. 3C, D).

**Figure 3.**
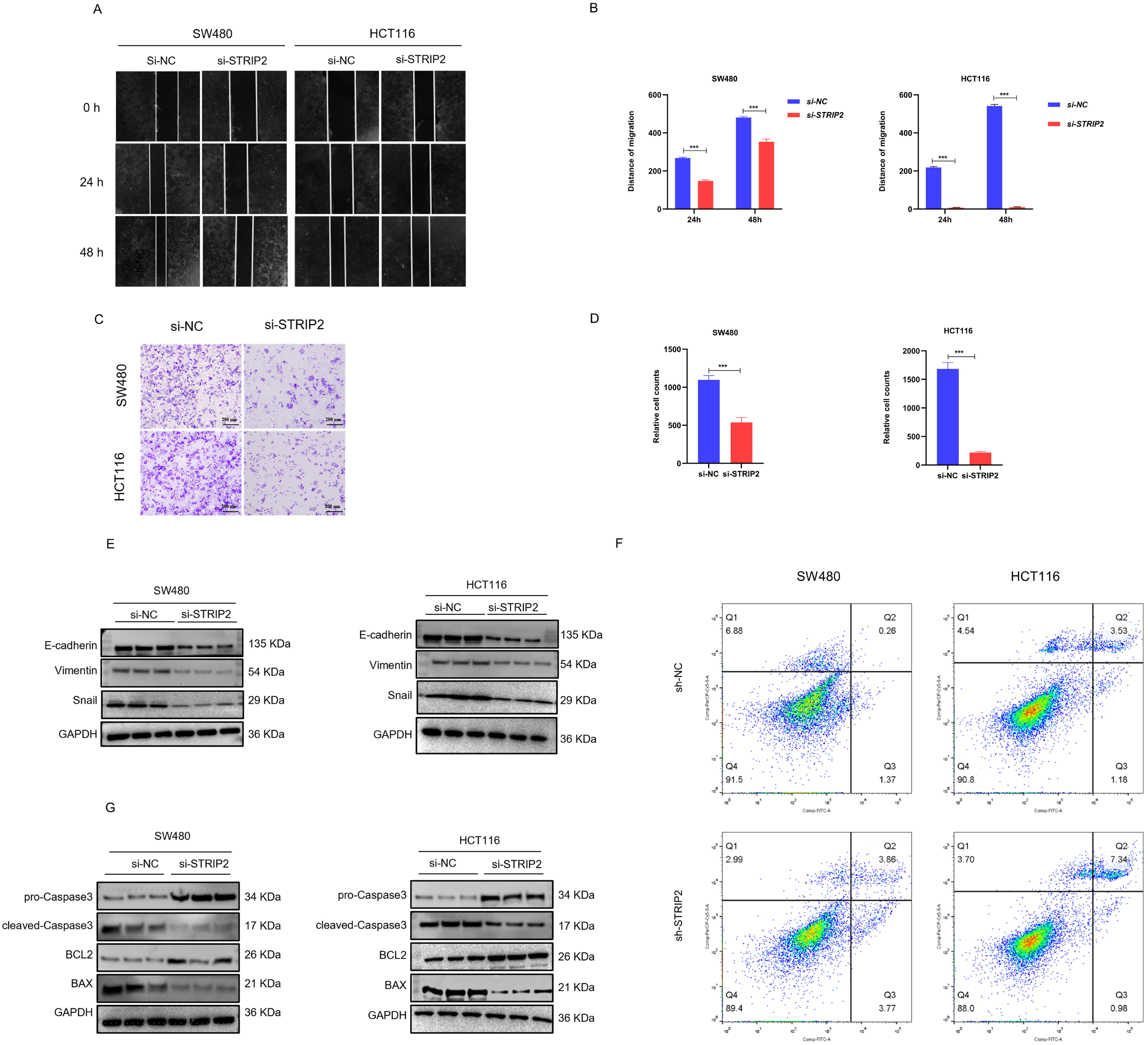
STRIP2 promotes CRC cell migration and invasion and inhibits CRC cell apoptosis. (A, B) Wound healing assays assessing migratory capacity in STRIP2-knockdown SW480 and HCT116 cells (n=3). (C, D) Transwell invasion assays in STRIP2-knockdown SW480 and HCT116 cells (n=3). (E) Western blot analysis of epithelial–mesenchymal transition (EMT) marker proteins in STRIP2-knockdown SW480 and HCT116 cells (n=3). (F) Flow cytometric quantification of apoptosis via Annexin V/PI staining in STRIP2-knockdown SW480 and HCT116 cells (n=3). (G) Western blot analysis of apoptosis-related protein levels (n=3) in STRIP2-knockdown SW480 and HCT116 cells (n=3). Data are presented as mean ± SD. *P <0.05, ** P<0.01, ***P<0.001. STRIP2: striatin-interacting phosphatase and kinase subunit 2; CRC: colorectal cancer; EMT: epithelial–mesenchymal transition; Annexin V/PI: Annexin V/propidium iodide.

At the molecular level, Western blot analysis of EMT markers demonstrated that STRIP2 knockdown upregulated the epithelial marker E-cadherin, while downregulating the mesenchymal marker vimentin and the core EMT transcription factor Snail (Fig. 3E). These results suggest that STRIP2 drives CRC cell migration and invasion by promoting EMT progression.

We subsequently examined the effect of STRIP2 on cell apoptosis. Flow cytometry analysis with Annexin V/PI double staining showed that the percentage of apoptotic cells was significantly increased in si-STRIP2 groups relative to si-NC groups in both SW480 and HCT116 cell lines (P < 0.05, Fig. 3F). Western blot analysis of apoptosis-related proteins further confirmed that STRIP2 knockdown elevated the levels of cleaved caspase-3 and the pro-apoptotic protein BAX, while reducing the levels of pro-caspase-3 and the anti-apoptotic protein BCL2 (Fig. 3G). These data indicate that STRIP2 suppresses CRC cell apoptosis by modulating the mitochondrial apoptosis pathway.

### STRIP2 promotes CRC tumor growth and metastasis in vivo

Firstly, we established subcutaneous xenograft models using HCT116 cells stably transduced with sh-STRIP2 or sh-NC. Compared with the control group, mice in the sh-STRIP2 group exhibited significantly slower tumor growth, smaller tumor volume and lower tumor weight (all P < 0.001, Fig. 4A-C). Histopathological examination revealed reduced mitotic activity in tumors from STRIP2 knockdown xenografts stained with H&E (Fig. 4D). IHC staining showed that the percentage of Ki-67+ cells indicating proliferative activity, was also lower in these tumors (Fig. 4D). ELISA results showed that CEA and CA199 levels were significantly lower in STRIP2 knockdown tumors compared to controls (both P < 0.01, Fig. 4E, F).

**Figure 4.**
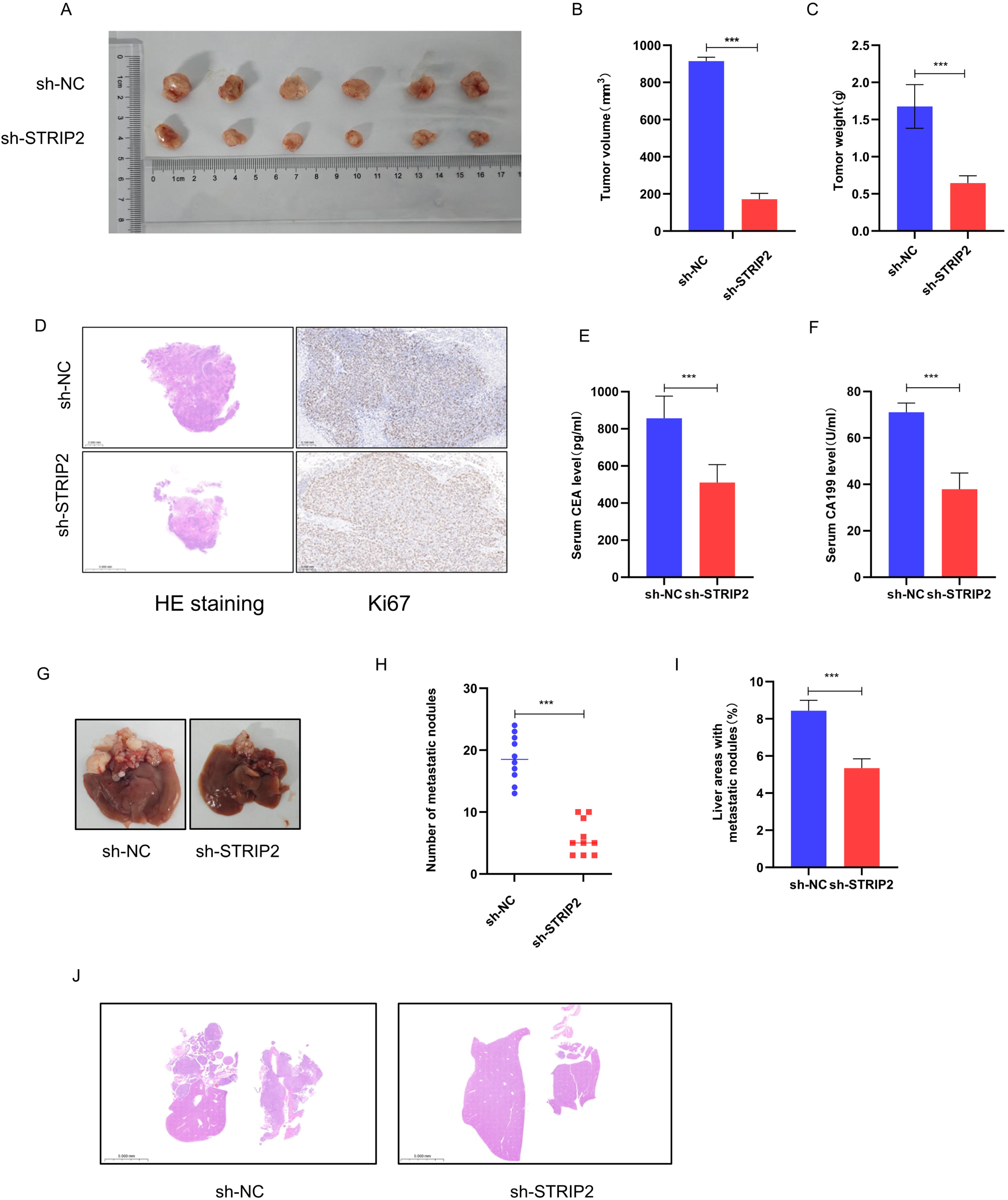
STRIP2 facilitates primary colorectal tumor growth and liver metastasis in vivo. (A –C) Subcutaneous xenograft model established by injecting HCT116 cells stably expressing control shRNA (sh-NC) or STRIP2-targeted shRNA (sh-STRIP2) into BALB/c nude mice (n=6): (A) Representative images of xenograft tumors, (B) Endpoint tumor volume, (C) Endpoint tumor weight. (D) Hematoxylin and eosin (H&E) and Ki-67 IHC staining of paraffin-embedded xenograft tumor sections. Scale bar, 50 μm. (E, F) ELISA detection of serum tumor markers CEA (E) and CA199 (F) in mice (n=6). (G–J) Liver metastasis model established via splenic injection (n=6): (G) Representative images of liver metastatic nodules, (H) Quantification of total metastatic nodule count, (I) Quantification of total metastatic lesion area, (J) H&E staining of liver metastatic tissues. Scale bars: 50 μm. Data are presented as mean ± SD. *P <0.05, ** P<0.01, ***P<0.001. STRIP2: striatin-interacting phosphatase and kinase subunit 2; shRNA: short hairpin RNA; BALB/c: BALB/c nude mouse strain; H&E: hematoxylin and eosin; IHC: immunohistochemistry; ELISA: enzyme-linked immunosorbent assay; CEA: carcinoembryonic antigen; CA199: carbohydrate antigen 19-9.

We further established a spleen injection-induced liver metastasis model to evaluate the impact of STRIP2 on CRC metastatic dissemination. Gross examination and H&E staining of liver tissues revealed that STRIP2 knockdown significantly reduced both the number and the area of metastatic nodules in the liver (all P < 0.001, Fig. 4G–J). These *in vivo* findings consistently support that STRIP2 promotes CRC tumor growth and metastatic progression.

### Transcriptomic profiling

To investigate the molecular mechanisms underlying STRIP2’s role in CRC progression, we performed transcriptomic profiling using RNA sequencing (RNA-seq) in stably transducted si-STRIP2 HCT116 cells vs si-NC cells.

As shown in the volcano plot (Fig. 5A), among a total of 32,799 detected transcripts, 764 genes were identified as significantly differentially expressed (adjusted P < 0.05, |log Fold Change| > 1) following STRIP2 depletion. Specifically, 262 genes were significantly upregulated, while 502 genes were significantly downregulated in si-STRIP2 cells relative to si-NC controls. The predominance of downregulated DEGs indicated that STRIP2 mainly acts as a positive regulator of gene expression programs in CRC cells, and its knockdown induces broad transcriptomic remodelling.

**Figure 5.**
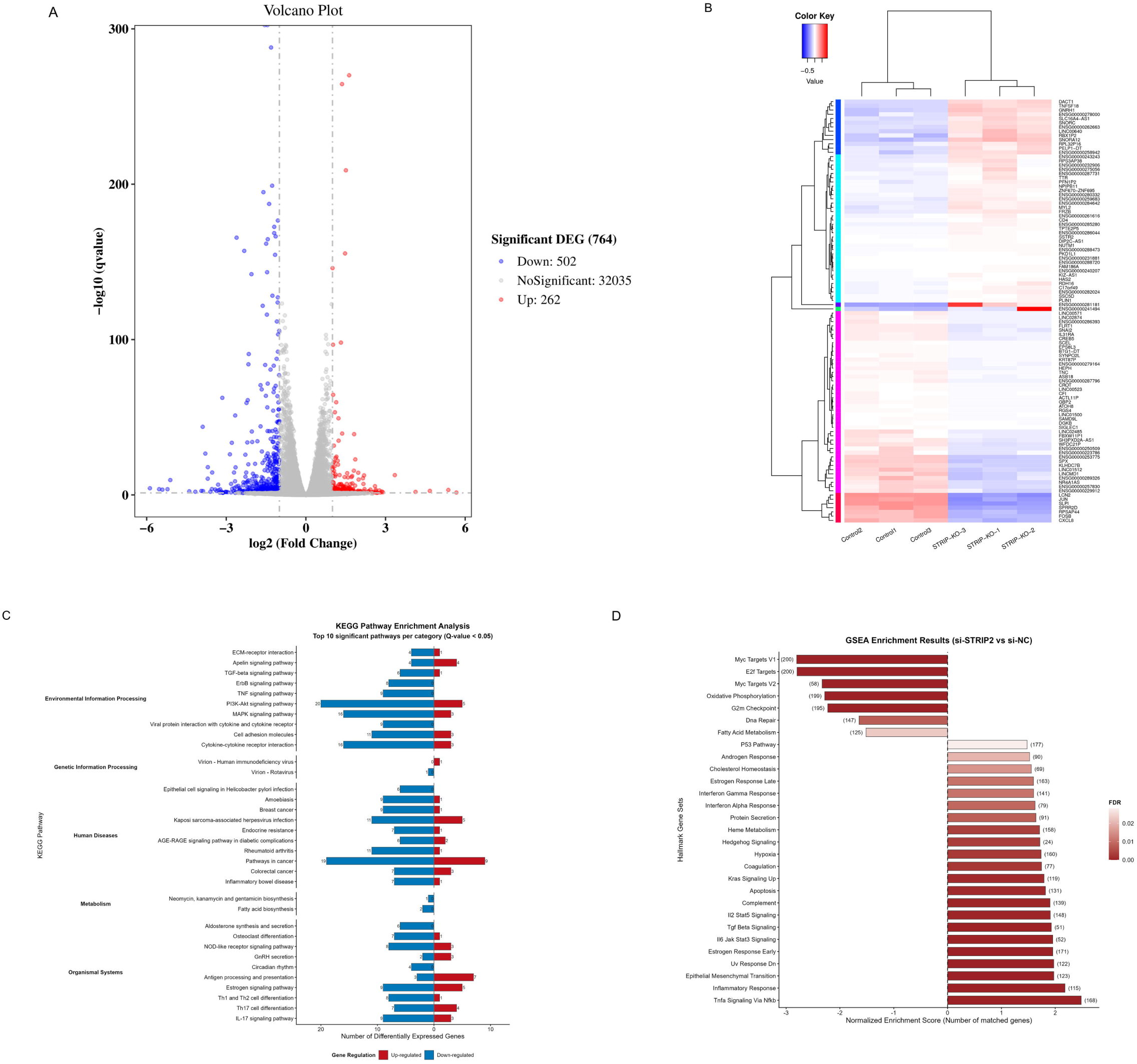
Comparative transcriptomic profiling identifies differentially expressed genes (DEGs) upon STRIP2 silencing in HCT116 CRC cells. (A) Volcano plot of DEGs between si-STRIP2 and si-NC HCT116 cell populations. (B) Heatmap of normalized DEG expression. (C) KEGG pathway enrichment of DEGs (|log FC|>1, P<0.05). (D) Gene set enrichment analysis (GSEA). Data are shown as mean ± SD; *P <0.05, ** P<0.01, ***P<0.001. STRIP2: striatin-interacting phosphatase and kinase subunit 2; CRC: colorectal cancer; DEGs: differentially expressed genes; si: silencing; NC: negative control; KEGG: Kyoto Encyclopedia of Genes and Genomes; log FC: log fold change; GSEA: gene set enrichment analysis.

Unsupervised hierarchical clustering of the top DEGs was visualized via a heatmap (Fig. 5B). DEGs were clearly partitioned into two major expression modules: genes with elevated expression in si-STRIP2 cells and genes with reduced expression in si-STRIP2 cells. Notably, LCN2, a well-characterized regulator of iron homeostasis and ferroptosis (24), was among the most prominently downregulated genes upon STRIP2 knockdown, which was selected for further mechanistic investigation.

KEGG pathway enrichment analysis results showed that DEGs were significantly enriched in multiple cancer-related signalling pathways (Fig. 5C). Among them, the IL-17 signalling pathway was one of the most significantly enriched pathways, suggesting that inflammatory signalling mediated by IL-17 is a key downstream cascade regulated by STRIP2. Other enriched pathways included the MAPK signalling pathway, pathways associated with cell differentiation, metabolic pathways, and human disease-related pathways. These enrichment results were consistent with the observed phenotypes of enhanced proliferation, migration and invasion driven by STRIP2 in CRC.

GSEA pathway enrichment analysis results showed that pathways positively associated with high STRIP2 expression (enriched in si-NC group) included EMT, TGF-beta signalling, Kras signalling up, estrogen response and inflammatory response and TNF-alpha signalling pathways (Fig. 5D). These pathways collectively contribute to cell motility, stromal remodelling, metastatic potential and inflammatory microenvironment formation, in line with the pro-metastatic function of STRIP2.

### STRIP2 stabilizes LCN2 protein through inhibiting its ubiquitin-proteasomal degradation

LCN2 was identified as a prominent downstream effector from transcriptomic data. We first validated that STRIP2 knockdown significantly reduced LCN2 protein levels in both SW480 and HCT116 cells (Fig. 6A). To determine whether STRIP2 regulates LCN2 at the post-translational level, we treated cells with the proteasome inhibitor MG132. As shown in Fig. 6B, MG132 treatment caused marked accumulation of LCN2 protein. It largely abolished the difference in LCN2 levels between control and oe-STRIP2 293T/17 cells, indicating that STRIP2 modulates LCN2 abundance via the ubiquitin-proteasome degradation pathway.

**Figure 6.**
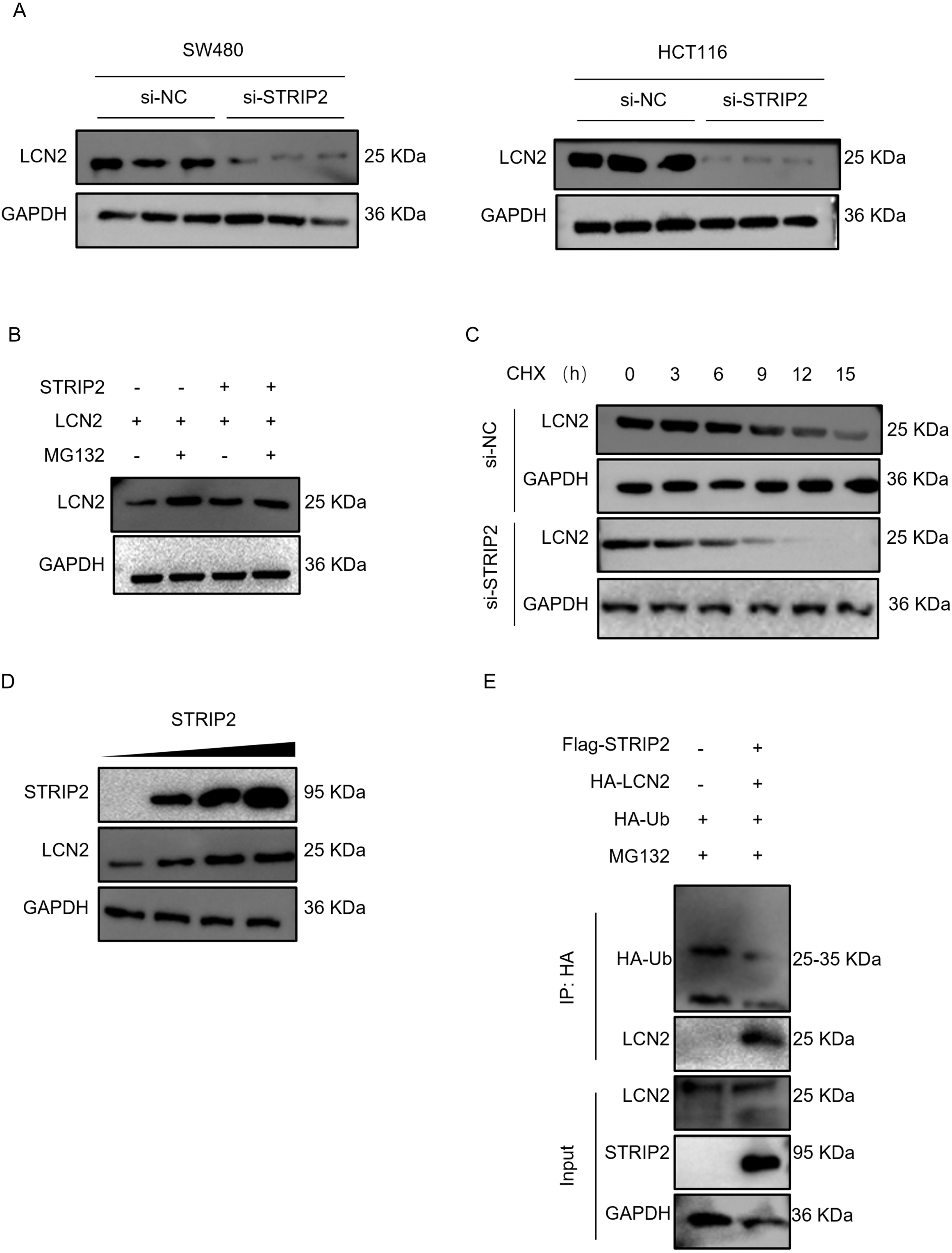
STRIP2 stabilizes LCN2 protein post-translationally by blocking ubiquitin-mediated proteasomal degradation. (A) Western blot of LCN2 in si-NC and si-STRIP2 SW480 and HCT116 cells. (B) Western blot of LCN2 and STRIP2 in 293T/17 cells with empty vector, single or combined LCN2/STRIP2 overexpression plasmids, with/without 6 h MG132 (20 μmol/l) treatment. (C) Time-course western blot measuring LCN2 half-life in si-NC and si-STRIP2 HCT116 cells treated with CHX (100 μg/ml) for 0–15 h. (D) Western blot detecting exogenous LCN2 and STRIP2 in co-transfected 293T/17 cells. (E) Co-IP assay verifying STRIP2-mediated inhibition of LCN2 ubiquitination. Data are shown as mean ± SD; ***P<0.001. STRIP2: striatin-interacting phosphatase and kinase subunit 2; LCN2: lipocalin 2; MG132: proteasome inhibitor; CHX: cycloheximide; co-IP: co-immunoprecipitation; 293T/17: human embryonic kidney cell line.

We further performed cycloheximide (CHX) chase assays to measure LCN2 protein half-life. STRIP2 knockdown significantly accelerated the degradation of endogenous LCN2 protein and shortened its half-life in HCT116 cells (Fig. 6C).

To directly examine the effect of STRIP2 on LCN2 ubiquitination, we conducted co-immunoprecipitation (Co-IP) assays in 293T/17 cells co-transfected with HA-LCN2, HA-Ub and Flag-STRIP2. The results showed that STRIP2 overexpression markedly decreased LCN2 polyubiquitination (Fig. 6D, E). Collectively, these data demonstrate that STRIP2 stabilizes LCN2 protein by inhibiting its polyubiquitination and subsequent proteasomal degradation.

### STRIP2 suppresses ferroptosis in CRC cells

Ferroptosis is an iron-dependent regulated cell death process frequently dysregulated in CRC. Our transcriptomic data showed LCN2 is the most prominently downregulated gene upon STRIP2 knockdown; given that LCN2 suppresses ferroptosis in CRC by upregulating ferroptosis-related genes (e.g., SLC7A11, GPX4) (24), we investigated whether STRIP2 affects ferroptotic cell death in CRC cells.

Western blot assays revealed that oe-STRIP2 decreased the level of the pro-ferroptotic protein ACSL4 (26, 27), while increasing the levels of ferroptosis suppressors SLC7A11(28), GPX4 (29), FSP1 (30, 31), and NRF2 (32); on the contrary, STRIP2 knockdown exerted the opposite effects (Fig. 7A).

**Figure 7.**
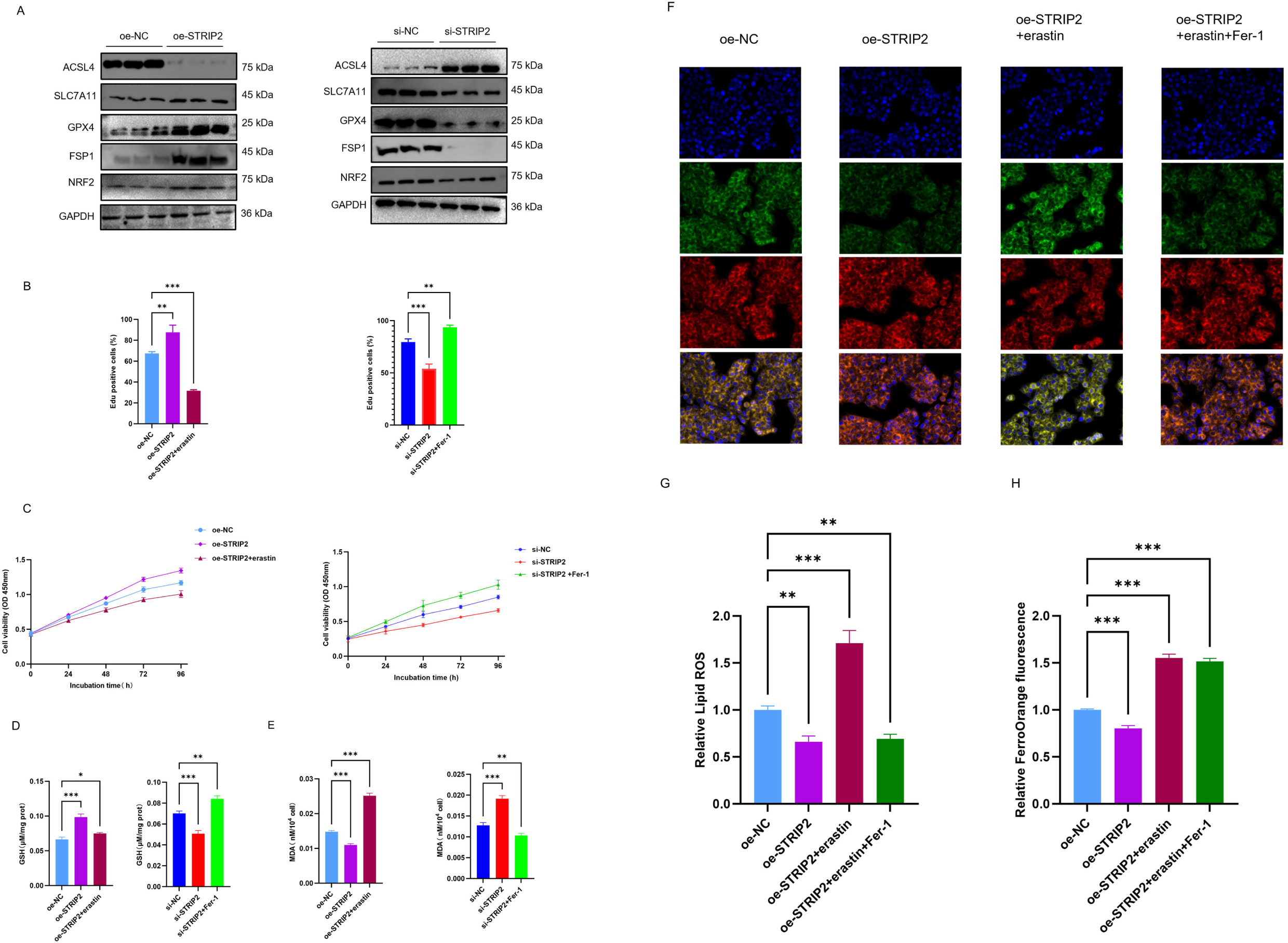
STRIP2 inhibits ferroptosis in CRC cells by regulating lipid peroxidation and iron homeostasis. (A) Western blot of core ferroptosis regulatory proteins (ACSL4, SLC7A11, GPX4, FSP1 and NRF2) regulators in oe-NC, oe-STRIP2, si-NC and si-STRIP2 HCT116 cells (n = 3). (B-E) Functional ferroptosis assays in oe-NC, oe-STRIP2 and oe-STRIP2 HCT116 cells + erastin (n = 6): and si-NC, si-STRIP2 and si-STRIP2 HCT116 cells + Fer-1 groups (n = 6): (B) EdU assay, (C) CCK-8 viability assay, (D) intracellular GSH and (E) intracellular MDA quantification. (F-G) Lipid ROS quantification in oe-NC, oe-STRIP2, oe-STRIP2+erastin, oe-STRIP2+erastin+Fer-1 (n = 3). (F) Representative immunofluorescence images of lipid ROS. Rows show DAPI (blue), lipid ROS green channel, lipid ROS red channel and merged images. (G) Quantitative lipid ROS fluorescence intensity. (H) Quantification of labile intracellular iron fluorescence intensity (n=3). Data are shown as mean ± SD. ***P<0.001. STRIP2: striatin-interacting phosphatase and kinase subunit 2; oe: overexpression; si: silencing; ACSL4, acyl-CoA synthetase long-chain family member 4; SLC7A11, solute carrier family 7 member 11; GPX4, glutathione peroxidase 4; FSP1, ferroptosis suppressor protein 1; NRF2, nuclear factor erythroid 2-related factor 2; erastin: ferroptosis inducer; Fer-1: ferrostatin-1 (ferroptosis inhibitor); GSH: reduced glutathione; MDA: malondialdehyde; lipid ROS: lipid reactive oxygen species; DAPI: 4’,6-diamidino-2-phenylindole.

We then performed experiments using the ferroptosis inducer erastin and the ferroptosis inhibitor Fer-1. Functionally, EdU (Fig. 7B) and CCK-8 (Fig. 7C) assays showed that oe-STRIP2 promoted CRC cell proliferation, while erastin treatment partially abrogated this pro-proliferative effect. Consistently, oe-STRIP2 overexpression increased intracellular GSH levels (Fig. 7D) and decreased MDA levels, whereas ferroptosis inducer erastin treatment reversed these metabolic changes (Fig. 7D, E). In contrast, si-STRIP2 inhibited cell proliferation, induced GSH depletion, and promoted MDA accumulation; Fer-1 treatment significantly reversed these effects (Fig. 7B-E).

Lipid peroxidation is a key feature of ferroptosis. We also used immunofluorescence to measure lipid ROS and intracellular labile iron in four colorectal cancer cell groups: oe-NC, oe-STRIP2, oe-STRIP2 + erastin, and oe-STRIP2 + erastin + Fer-1. DAPI (blue) stained nuclei for normalization, green signals denoted lipid ROS, and red signals represented labile iron, with merged images confirming cytoplasmic co-localization of the two ferroptotic markers (Fig. 7F).

Basal lipid ROS and intracellular labile iron levels marginally increased in oe-STRIP2 cells relative to oe-NC controls. Erastin treatment robustly elevated lipid ROS and labile iron in oe-STRIP2 cells, as shown by stronger fluorescence intensity and statistically significant quantitative bar plots (Fig. 7F, G). Concurrent Fer-1 supplementation reversed erastin-mediated lipid ROS overaccumulation (Fig. F, G), but did not significantly reverse iron overload (Fig. H). These findings suggest that STRIP2 acts as a positive regulator of ferroptosis by modulating core ferroptosis effector proteins, and that the pro-ferroptotic effect conferred by STRIP2 overexpression is reversible with Fer-1 intervention.

Taken together, these results indicate that STRIP2 inhibits ferroptosis in CRC cells, thereby enhancing cell survival and proliferative capacity. Collectively, these results suggest that STRIP2 regulates ferroptosis by stabilizing LCN2 expression and preventing its proteasomal degradation. STRIP2-mediated LCN2 stabilization enhances anti-ferroptotic effects, promoting CRC cell survival and providing a novel link between STRIP2 and ferroptosis regulation in CRC.

## Discussion

In the present study, we demonstrated that STRIP2 was significantly upregulated in CRC tissues and cell lines. This upregulation pattern aligns with that observed in other malignancies, including non-small cell lung cancer (12), hepatocellular carcinoma (33), and breast cancer (34), suggesting STRIP2’s potential as a pan-cancer diagnostic biomarker.

Functionally, STRIP2 knockdown suppressed CRC cell proliferation, migration, invasion, and EMT, while inducing mitochondrial apoptosis *in vitro*. Additionally, it inhibited subcutaneous tumor growth and liver metastasis *in vivo*. Furthermore, we identified a novel post-translational regulatory mechanism by which STRIP2 stabilizes the LCN2 protein by inhibiting its polyubiquitination and subsequent proteasomal degradation, thereby conferring ferroptosis resistance in CRC cells. Although LCN2 has long been recognized as an oncogenic effector that modulates iron homeostasis and lipid peroxidation in CRC, the upstream regulators governing its protein stability through the ubiquitin-proteasome system remain elusive (35) and warrant further investigation.

In addition, while existing research on STRIP2 has predominantly focused on its role in PI3K/AKT/mTOR (11) and Hippo (36) signaling pathways, our discovery of the STRIP2-LCN2-ferroptosis axis expands its functional repertoire to include regulation of ferroptosis and metabolic reprogramming. This adds a new layer of mechanistic understanding to STRIP2’s oncogenic activity.

These findings extend the current understanding of STRIP2 in cancer biology, providing a novel molecular framework for CRC pathogenesis, positioning the STRIP2-LCN2 axis as a promising therapeutic target for CRC intervention.

### Molecular mechanism of STRIP2 stabilizing LCN2 to suppress ferroptosis in CRC

Ubiquitination is a crucial post-translational modification that regulates the turnover of ferroptosis-related proteins (37). Several deubiquitinases, including USP8 and OTUB1, have been shown to stabilize key anti-ferroptotic proteins, such as GPX4 and SLC7A11, thereby conferring resistance to ferroptosis in CRC (38). Conversely, E3 ubiquitin ligases promote substrate degradation, thereby making tumor cells more sensitive to ferroptosis (39).

In our study, we employed CHX and Co-IP assays to identify STRIP2 as a novel upstream stabilizer of LCN2. This finding complemented the existing regulatory mechanisms of LCN2, which are primarily controlled at the transcriptional level by IL-17/NRF2 signaling (40).

Once stabilized by STRIP2, accumulated LCN2 exerts dual anti-ferroptotic functions through the three primary branches of the ferroptosis machinery. In the iron metabolism pathway, LCN2 acts as an iron chelator, reducing the intracellular labile iron pool (41). This action mitigates free radical production from the Fenton reaction and prevents the initiation of membrane lipid peroxidation (42). Along the cysteine-GSH-GPX4 antioxidant axis, increased LCN2 levels transcriptionally activate SLC7A11, enhancing extracellular cystine uptake, supporting intracellular GSH biosynthesis, and preserving GPX4 enzymatic activity to clear toxic lipid peroxides (43). Additionally, LCN2 upregulates FSP1 and NRF2, establishing a GPX4-independent redundant antioxidant network (44).

In lipid metabolism regulation, LCN2 significantly limits ACSL4 expression, thereby reducing the incorporation of polyunsaturated fatty acids into cell membrane phospholipids and limiting the substrates available for lipid peroxidation (45, 46). Supporting these biochemical changes, we show that STRIP2 overexpression increases intracellular GSH concentrations and decreases MDA levels (a marker of lipid peroxidation), alongside elevated SLC7A11, GPX4, FSP1, and NRF2, and reduced ACSL4 levels (47). STRIP2 knockdown downregulates LCN2, collapses the multi-layered anti-ferroptotic defence, decreases GSH levels, and increases MDA content, ultimately restoring ferroptosis sensitivity in CRC cells. Using the ferroptosis inducer erastin and the inhibitor Fer-1 further confirms that ferroptosis is the specific downstream effector driving STRIP2-influenced CRC proliferation.

In summary, STRIP2-mediated LCN2 stabilization allows CRC cells to evade ferroptotic elimination under oxidative stress within a nutrient-poor tumor microenvironment. This finding highlights STRIP2’s pronounced oncogenic role in promoting primary tumor growth and distant liver metastasis.

Consistent with previous studies showing that IL-17 cytokines transcriptionally activate LCN2 expression in inflammation-associated CRC (48), our research adds an important post-translational regulatory layer governed by STRIP2. This establishes a dual transcriptional and post-translational regulatory network that sustains persistent LCN2 accumulation and robust ferroptosis resistance in IL-17-rich inflammatory CRC microenvironments. Furthermore, LCN2 facilitates CRC invasion by promoting focal adhesion formation and c-Src activation (49), providing further insight into the pro-metastatic phenotype induced by the STRIP2-LCN2 axis.

### Translational clinical implications of STRIP2-LCN2-ferroptosis axis for CRC treatment

The STRIP2-LCN2-ferroptosis regulatory cascade offers promising clinical opportunities for treating refractory CRC through targeted therapies. STRIP2 and LCN2 expression levels in CRC tissues can serve as predictive biomarkers for ferroptosis-inducing combination therapies. Patients with high levels of these markers typically exhibit enhanced anti-ferroptotic capacity and poor responses to conventional chemotherapy, making them ideal candidates for STRIP2 inhibitors in combination with ferroptosis inducers (50). Conversely, patients with low expression may benefit more from standard chemotherapy.

Inhibiting STRIP2 provides dual anti-tumor benefits. It attenuates the pro-tumor IL-17 inflammatory response and also promotes LCN2 degradation, thereby weakening the anti-ferroptotic defense and rendering resistant CRC cells susceptible to ferroptosis (51). The resulting lipid peroxidation can release damage-associated molecular patterns (DAMPs), thereby remodeling the immunosuppressive tumor microenvironment (TME), and enhancing CD8 T cell infiltration, synergizing with anti-PD-1 immunotherapy (52). For chemo-refractory CRC with high STRIP2/LCN2, dual targeting of STRIP2 and ferroptosis inducers represents a novel synthetic lethal strategy, as these tumors are highly dependent on LCN2-mediated ferroptosis evasion for survival. Combining a STRIP2 inhibitor, a ferroptosis inducer, and an immune checkpoint blocker could represent a novel treatment strategy for advanced chemotherapy-resistant CRC (53), expanding the translational potential beyond chemotherapy.

### Study limitations and future research

Limitations of our study include the small sample size (5 pairs of CRC and adjacent non-tumor tissues), which restricts our ability to draw meaningful clinical correlations. Future research should validate STRIP2 expression in larger cohorts and include multivariate analyses to assess its prognostic value. This study is also limited by the use of subcutaneous xenograft models, which do not fully recapitulate the intestinal inflammatory microenvironment or liver metastasis. Future studies can use orthotopic CRC and ApcMin/+ mice to confirm STRIP2-mediated ferroptosis regulation in situ and evaluate STRIP2 inhibitors combined with ferroptosis inducers in patient-derived xenograft (PDX) models for clinical translation. Finally, the long-term effects of STRIP2 knockdown on tumor dynamics warrant further investigation. Future studies should also focus on developing specific STRIP2 inhibitors and evaluating their synergistic effects with existing drug treatments.

In conclusion, this study highlights STRIP2’s role in CRC progression, demonstrating its significant upregulation in CRC tissues and cell lines. STRIP2 knockdown slows CRC growth, migration, and liver metastasis, and triggers tumor cell apoptosis.

Transcriptome sequencing revealed that STRIP2 depletion downregulated LCN2. STRIP2 interacts with LCN2, preventing its degradation and enhancing anti-ferroptotic effects via the SLC7A11/GSH/GPX4 antioxidant axis, while suppressing ACSL4 expression, thereby limiting lipid peroxidation.

The STRIP2-LCN2-ferroptosis signaling axis reveals a dual transcriptional-post-translational regulatory network that sustains persistent ferroptosis resistance in inflammatory CRC, presenting a potential therapeutic target for combined treatment strategies for advanced chemo-resistant CRC.

## Supporting information

Western blot raw images

## Acknowledgments

We thank our colleagues for their support in this study.

## Availability of data and materials

The raw RNA sequencing data generated in this study have been deposited into the Gene Expression Omnibus Repository under accession number GSE333323. Other data are available from the corresponding author upon request.

## Authors’ contributions

CH, XY, SZ, XC, HH conducted investigations and data analysis. XY, HH, JD and WT conceived and designed the study. HH, XC and JD developed the methodology. XY acquired the research funding, and WT administered the whole research project. CH, XY, SZ and XC drafted the original manuscript; all authors reviewed and edited the manuscript. CH, XY, XC and HH verified the authenticity of all raw data. All authors have read and approved the final manuscript.

## Ethics approval and consent to participate

Human ethics approval for the use of human specimens was granted by the Committees for Ethical Review at Affiliated Jinhua Hospital, Zhejiang University School of Medicine (Jinhua, China; reference number: 20252220101), with written informed consent from all patients in accordance with the Declaration of Helsinki. All experimental mice used for this study were cared for under a protocol approved by the Experimental Animal Ethics Committee of Jinhua Institute of Food and Drug Inspection and Testing (reference number: AL-JSYJ202473) and were strictly adhered to the ARRIVE 2.0 guidelines.

## Patient consent for publication

Written informed consent forms were obtained from all patients for publication.

## Use of artificial intelligence tools

During the preparation of this work, artificial intelligence tools were used to improve the readability and language of the manuscript. The authors then revised and edited the content produced by the artificial intelligence tools as necessary and take full responsibility for the final content of the present manuscript.

## Abbreviations

ACSL4: acyl-CoA synthetase long-chain family member 4
BCA: bicinchoninic acid
CA199: carbohydrate antigen 199
CCK8: cell counting Kit 8
CEA: carcinoembryonic antigen
CHX: cycloheximide
COAD: colon adenocarcinoma
CRC: colorectal cancer
DAB: 3,3-diaminobenzidine
DAMPs: damage-associated molecular patterns
DEGs: differentially expressed genes
DMEM: Dulbecco’s modified Eagle’s medium
EdU: 5-ethynyl-2’-deoxyuridine
ELISA: enzyme-linked immunosorbent assay
EMT: epithelial-mesenchymal transition
FPKM: fragments per kilobase per million reads
GAPDH: glyceraldehyde-3-phosphate dehydrogenase
GEO: Gene Expression Omnibus
GPX4: glutathione peroxidase 4
GSEA: gene set enrichment analysis
GSH: glutathione
H&E: hematoxylin and eosin
HRP: horseradish peroxidase
IHC: immunohistochemistry
IL-17: interleukin-17
IP: immunoprecipitation
KEGG: Kyoto encyclopedia of genes and genomes
LCN2: lipocalin-2
limma: linear models for microarray data
MDA: malondialdehyde
MEM: minimal essential medium
MSigDB: molecular signatures database
NSCLC: non-small cell lung cancer
PI: propidium iodide
PMSF: phenylmethanesulfonylfluoride
PVDF: polyvinylidene difluoride
READ: rectal adenocarcinoma
RIPA: radioimmunoprecipitation assay
ROS: reactive oxygen species
SD: standard deviation
SDS: sodium dodecyl sulfate
SLC7A11: solute carrier family 7 member 11
STRIP2: Striatin-interacting protein 2
STRIPAK: striatum-interacting phosphatase and kinase
TBST: tris-buffered saline with Tween 20
TCGA: Cancer Genome Atlas
TME: tumor microenvironment

---

**Supplementary Table 1.**
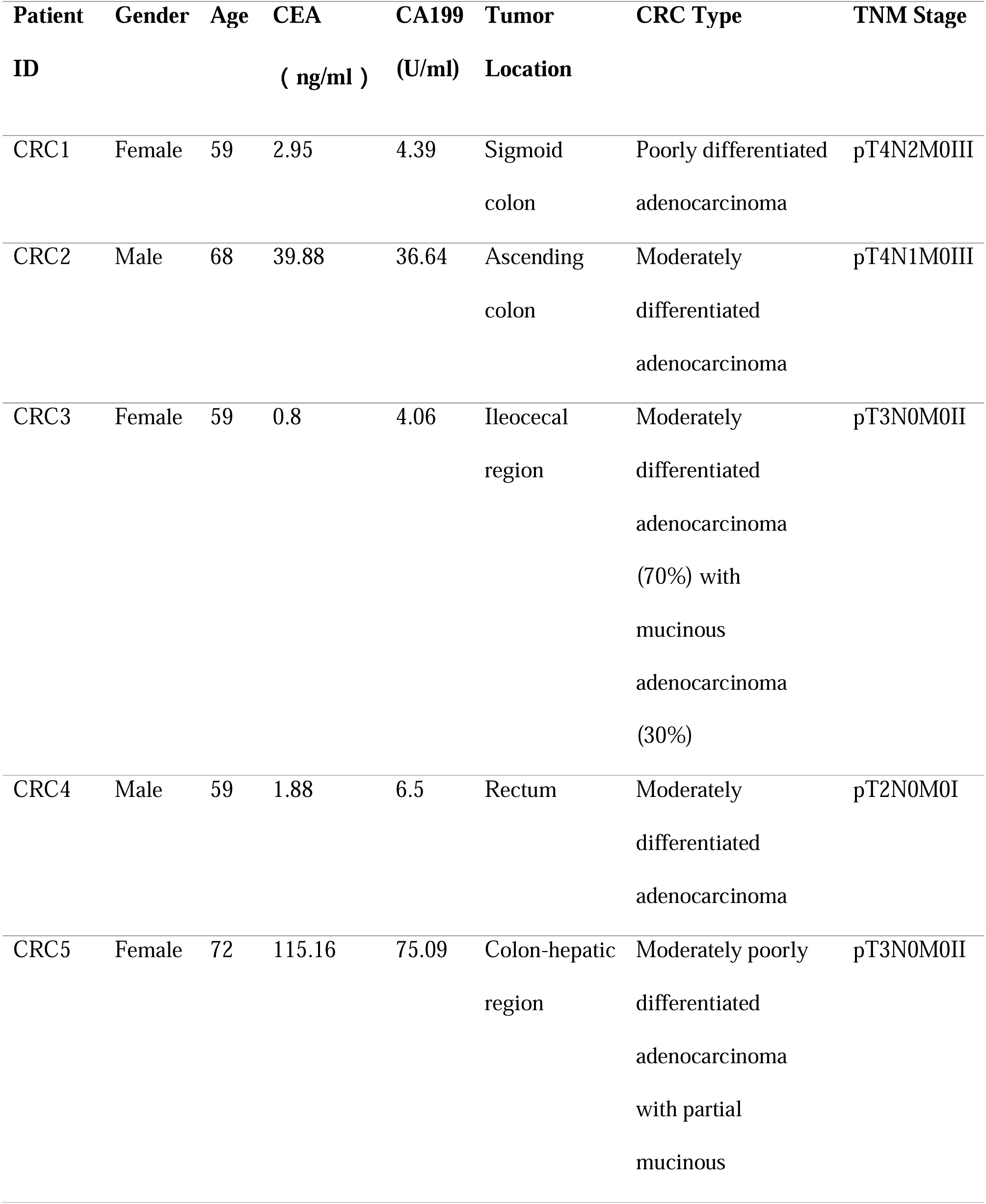
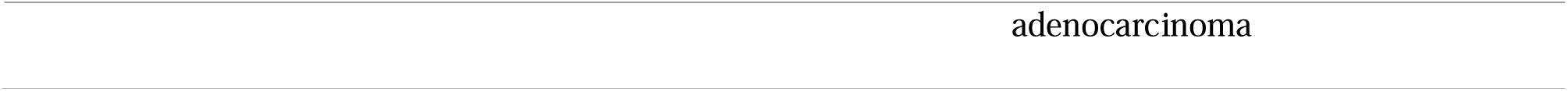
Patients’ clinical information.

| **Patient ID** | **Gender** | **Age** | **CEA**<br>**( ng/ml )** | **CA199**<br>**(U/ml)** | **Tumor Location** | **CRC Type** | **TNM Stage** |
| --- | --- | --- | --- | --- | --- | --- | --- |
| CRC1 | Female | 59 | 2.95 | 4.39 | Sigmoid colon | Poorly differentiated adenocarcinoma | pT4N2M0III |
| CRC2 | Male | 68 | 39.88 | 36.64 | Ascending colon | Moderately differentiated adenocarcinoma | pT4N1M0III |
| CRC3 | Female | 59 | 0.8 | 4.06 | Ileocecal region | Moderately differentiated adenocarcinoma (70%) with mucinous adenocarcinoma (30%) | pT3N0M0II |
| CRC4 | Male | 59 | 1.88 | 6.5 | Rectum | Moderately differentiated adenocarcinoma | pT2N0M0I |
| CRC5 | Female | 72 | 115.16 | 75.09 | Colon-hepatic region | Moderately poorly differentiated adenocarcinoma with partial mucinous | pT3N0M0II |

**Supplementary Table 2.**
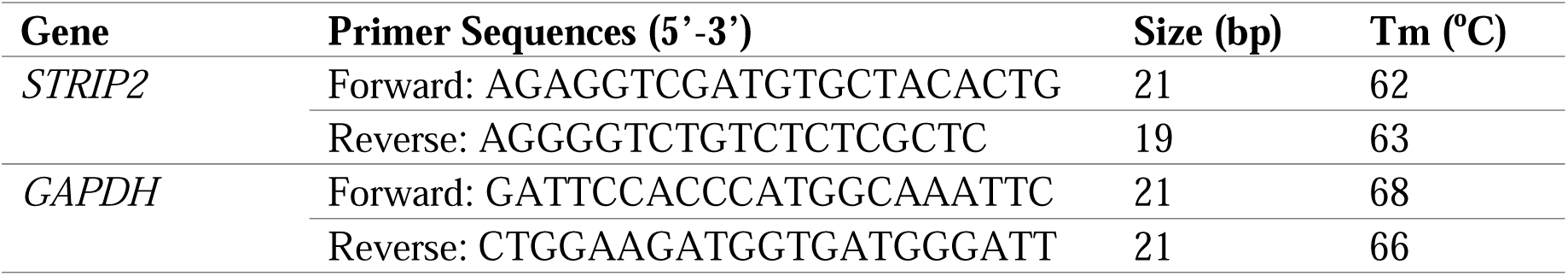
List of primers used in this study.

**Supplementary Table 3.** List of antibodies and dilutions used in this study.

| Antibody | Catalogue number<sup>#</sup> | Dilution factor |
| --- | --- | --- |
| STRIP2 | 25163-1-AP | 1: 1000 |
| GAPDH | 60004-1-Ig | 1: 50000 |
| E-cadherin | 20874-1-AP | 1: 5000 |
| Vimentin | 10366-1-AP | 1: 5000 |
| Snail | 12129-1-AP | 1: 1000 |
| Caspase-3 | 40924 | 1: 1000 |
| BCL2 | 60178-1-Ig | 1: 5000 |
| BAX | 50599-2-Ig | 1: 5000 |
| ACSL4 | 22401-1-AP | 1: 1000 |
| SLC7A11 | 26864-1-AP | 1: 1000 |
| GPX4 | 67763-1-Ig | 1: 2000 |
| FSP1 | 20886-1-AP | 1: 1000 |
| LCN2 | 83853-2-RR | 1: 1000 |
| NRF2 | 16396-1-AP | 1: 1000 |
| Ki-67 | 28074-1-AP | 1: 4000 |

<sup>#</sup> All primary antibodies are obtained from Proteintech, Wuhan, China.

## References

1. Li Q, Geng S, Luo H, Wang W, Mo YQ, Luo Q, Wang L, Song GB, Sheng JP and Xu B: Signaling pathways involved in colorectal cancer: pathogenesis and targeted therapy. Signal Transduct Target Ther 9: 266, 2024.

2. Heisser T, Hoffmeister M, Tillmanns H and Brenner H: Impact of demographic changes and screening colonoscopy on long-term projection of incident colorectal cancer cases in Germany: A modelling study. Lancet Reg Health Eur 20: 100451, 2022.

3. Xi Y and Xu P: Global colorectal cancer burden in 2020 and projections to 2040. Transl Oncol 14: 101174, 2021.

4. Soerjomataram I and Bray F: Planning for tomorrow: global cancer incidence and the role of prevention 2020-2070. Nat Rev Clin Oncol 18: 663–672, 2021.

5. Li Y, Bi Y, Li W, Piao Y, Piao J, Wang T and Ren X: Research progress on ferroptosis in colorectal cancer. Front Immunol 15: 1462505, 2024.

6. Patel SG, Karlitz JJ, Yen T, Lieu CH and Boland CR: The rising tide of early-onset colorectal cancer: a comprehensive review of epidemiology, clinical features, biology, risk factors, prevention, and early detection. Lancet Gastroenterol Hepatol 7: 262–274, 2022.

7. Shah SC and Itzkowitz SH: Colorectal Cancer in Inflammatory Bowel Disease: Mechanisms and Management. Gastroenterology 162: 715–730 e713, 2022.

8. Goudreault M, D’Ambrosio LM, Kean MJ, Mullin MJ, Larsen BG, Sanchez A, Chaudhry S, Chen GI, Sicheri F, Nesvizhskii AI, et al: A PP2A phosphatase high density interaction network identifies a novel striatin-interacting phosphatase and kinase complex linked to the cerebral cavernous malformation 3 (CCM3) protein. Mol Cell Proteomics 8: 157–171, 2009.

9. Chen A, Liu N, Xu C, Wu S, Liu C, Qi H, Ren Y, Han X, Yang K, Liu X, et al: The STRIPAK complex orchestrates cell wall integrity signalling to govern the fungal development and virulence of Fusarium graminearum. Mol Plant Pathol 24: 1139–1153, 2023.

10. Hwang J and Pallas DC: STRIPAK complexes: structure, biological function, and involvement in human diseases. Int J Biochem Cell Biol 47: 118–148, 2014.

11. Pan J, Zhang Y, He L, Wu Y, Xiao W, Zhang J and Xu Y: STRIP2 is regulated by the transcription factor Sp1 and promotes lung adenocarcinoma progression via activating the PI3K/AKT/mTOR/MYC signaling pathway. Genomics 116: 110923, 2024.

12. Zhang X, Chen Q, He Y, Shi Q, Yin C, Xie Y, Yu H, Bao Y, Wang X, Tang C, et al: STRIP2 motivates non-small cell lung cancer progression by modulating the TMBIM6 stability through IGF2BP3 dependent. J Exp Clin Cancer Res 42: 19, 2023.

13. Wu J, Lu G, Zhou S, Jin Z and Fang F: MicroRNA-30c-2-3p targets STRIP2 to suppress malignant progression of gastric cancer cells. J Biochem 171: 451–457, 2022.

14. Dai L, Zhou J, Li T, Qian Y, Jin L, Zhu C and Li S: STRIP2 silencing inhibits vascular smooth muscle cell proliferation and migration via P38-AKT-MMP-2 signaling pathway. J Cell Physiol 234: 22463–22476, 2019.

15. Srinivasan SP, Nemade H, Cherianidou A, Peng L, Cruz-Molina S, Rada-Iglesias A and Sachinidis A: Epigenetic mechanisms of Strip2 in differentiation of pluripotent stem cells. Cell Death Discov 8: 447, 2022.

16. Jaradat JH, Masharqa G, Al-Shnaikat RG, Mashal M, Al-Nusairi AM and Saeed A: The Multifaceted Role of Striatin-interacting Protein 2 (STRIP2) in Disease Pathogenesis and Cancer Progression. Anticancer Res 44: 5169–5174, 2024.

17. Dixon SJ and Olzmann JA: The cell biology of ferroptosis. Nature Reviews Molecular Cell Biology 25: 424–442, 2024.

18. Fischer K, Thewes L, Prozorovski T, Bayer M, Dietrich M, Lowin T, Albrecht P, Hartung HP, Meuth SG, Aktas O, et al: Fumarate-based drugs protect against neuroinflammation via upregulation of anti-ferroptotic pathways. J Neuroinflammation 22: 241, 2025.

19. Jiang C, Yan Y, Long T, Xu J, Chang C, Kang M, Wang X, Chen Y and Qiu J: Ferroptosis: a potential therapeutic target in cardio-cerebrovascular diseases. Mol Cell Biochem 480: 4379–4399, 2025.

20. Jiang Y, Zhang M and Sun M: ACSL4 at the helm of the lipid peroxidation ship: a deep-sea exploration towards ferroptosis. Front Pharmacol 16: 1594419, 2025.

21. Liu W, Xie X, Zong H, Li Y, Ding Y, Liu Z, Wan B, Xiao T, Lv F, Tang C, et al: Design, synthesis and biological evaluation of triazolothiadiazole derivatives as FSP1 inhibitors for sensitizing cancer cells to ferroptosis. Eur J Med Chem 293: 117737, 2025.

22. Liu MY, Li HM, Wang XY, Xia R, Li X, Ma YJ, Wang M and Zhang HS: TIGAR drives colorectal cancer ferroptosis resistance through ROS/AMPK/SCD1 pathway. Free Radic Biol Med 182: 219–231, 2022.

23. Lu H, Tong W, Jiang M, Liu H, Meng C, Wang K and Mu X: Mitochondria-Targeted Multifunctional Nanoprodrugs by Inhibiting Metabolic Reprogramming for Combating Cisplatin-Resistant Lung Cancer. ACS Nano 18: 21156–21170, 2024.

24. Feng M, Wu X, Zhang J, Chen P, Qian S and Chang C: Loss of Lipocalin2 confers cisplatin vulnerability through modulating NF-kB mediated ferroptosis via ferroportin. Am J Cancer Res 14: 2088–2102, 2024.

25. Chen Y, Li B, Quan J, Li Z, Li Y and Tang Y: Inhibition of Ferroptosis by Mesenchymal Stem Cell-Derived Exosomes in Acute Spinal Cord Injury: Role of Nrf2/GCH1/BH4 Axis. Neurospine 21: 642–655, 2024.

26. Doll S, Proneth B, Tyurina YY, Panzilius E, Kobayashi S, Ingold I, Irmler M, Beckers J, Aichler M, Walch A, et al: ACSL4 dictates ferroptosis sensitivity by shaping cellular lipid composition. Nat Chem Biol 13: 91–98, 2017.

27. Shao S, Li W, Hong Y, Zeng R, Zhu L, Yi L, Li Y, Wang Y, Huang H, Jiang X, et al: ZDHHC2-Dependent Palmitoylation Dictates Ferroptosis and Castration Sensitivity in Prostate Cancer via Controlling ACSL4 Degradation and Lipid Peroxidation. Adv Sci (Weinh) e14077, 2025.

28. Badgley MA, Kremer DM, Maurer HC, DelGiorno KE, Lee HJ, Purohit V, Sagalovskiy IR, Ma A, Kapilian J, Firl CEM, et al: Cysteine depletion induces pancreatic tumor ferroptosis in mice. Science 368: 85–89, 2020.

29. Yang WS, SriRamaratnam R, Welsch ME, Shimada K, Skouta R, Viswanathan VS, Cheah JH, Clemons PA, Shamji AF, Clish CB, et al: Regulation of ferroptotic cancer cell death by GPX4. Cell 156: 317–331, 2014.

30. Doll S, Freitas FP, Shah R, Aldrovandi M, da Silva MC, Ingold I, Goya Grocin A, Xavier da Silva TN, Panzilius E, Scheel CH, et al: FSP1 is a glutathione-independent ferroptosis suppressor. Nature 575: 693–698, 2019.

31. Bersuker K, Hendricks JM, Li Z, Magtanong L, Ford B, Tang PH, Roberts MA, Tong B, Maimone TJ, Zoncu R, et al: The CoQ oxidoreductase FSP1 acts parallel to GPX4 to inhibit ferroptosis. Nature 575: 688–692, 2019.

32. Dodson M, Castro-Portuguez R and Zhang DD: NRF2 plays a critical role in mitigating lipid peroxidation and ferroptosis. Redox Biol 23: 101107, 2019.

33. Yang W, Chen Y, Guo H, Zhang J, Wang S and Peng L: STRIP2 promotes hepatocellular carcinoma progression and immune evasion: a potential prognostic biomarker and therapeutic target. J Gastrointest Oncol 16: 2314–2335, 2025.

34. Li AX, Zeng JJ, Martin TA, Ye L, Ruge F, Sanders AJ, Khan E, Dou QP, Davies E and Jiang WG: Striatins and STRIPAK complex partners in clinical outcomes of patients with breast cancer and responses to drug treatment. Chin J Cancer Res 35: 365–385, 2023.

35. Liu X, Liu J, Chen W, Zhang J, Wei Y and Sun L: Ubiquitin regulatory code: unraveling tumor cell ferroptosis. Crit Rev Oncol Hematol 220: 105172, 2026.

36. Lu M and Luo X: Structural Studies of Core Hippo Pathway Components. Cold Spring Harb Perspect Biol 2026.

37. Meng Y, Sun H, Li Y, Zhao S, Su J, Zeng F, Deng G and Chen X: Targeting Ferroptosis by Ubiquitin System Enzymes: A Potential Therapeutic Strategy in Cancer. Int J Biol Sci 18: 5475–5488, 2022.

38. Wang Y, Liu B, Zhuang Y, Zhang Y, Dan W, Ding P, Wei Y, Guo C, Li M, Wang C, et al: USP20 governs tyrosine kinase inhibitors resistance through ferroptosis evasion by targeting GPX4 in cancers. Redox Biol 92: 104086, 2026.

39. Nguyen KT, Mun SH, Yang J, Lee J, Seok OH, Kim E, Kim D, An SY, Seo DY, Suh JY, et al: The MARCHF6 E3 ubiquitin ligase acts as an NADPH sensor for the regulation of ferroptosis. Nat Cell Biol 24: 1239–1251, 2022.

40. Ferreira MC, Whibley N, Mamo AJ, Siebenlist U, Chan YR and Gaffen SL: Interleukin-17-induced protein lipocalin 2 is dispensable for immunity to oral candidiasis. Infect Immun 82: 1030–1035, 2014.

41. Che R, Wang Q, Li M, Shen J and Ji J: Quantitative Proteomics of Tissue-Infiltrating T Cells From CRC Patients Identified Lipocalin-2 Induces T-Cell Apoptosis and Promotes Tumor Cell Proliferation by Iron Efflux. Mol Cell Proteomics 23: 100691, 2024.

42. Noguchi N, Saito Y and Niki E: Lipid peroxidation, ferroptosis, and antioxidants. Free Radic Biol Med 237: 228–238, 2025.

43. Wientjens C, Doverman M, Zurkovic J, More T, Surendar J, Nesic S, Sarici C, Utecht TD, Pohl J, Pollock J, et al: Tolerance to ferroptosis facilitates lipid metabolism and pathogenic type 2 immunity in allergic airway inflammation. Immunity 59: 322–338 e329, 2026.

44. Liu J, Pang SY, Zhou SY, He QY, Zhao RY, Qu Y, Yang Y and Guo ZN: Lipocalin-2 aggravates blood-brain barrier dysfunction after intravenous thrombolysis by promoting endothelial cell ferroptosis via regulating the HMGB1/Nrf2/HO-1 pathway. Redox Biol 76: 103342, 2024.

45. Deng L, He S, Li Y, Ding R, Li X, Guo N and Luo L: Identification of Lipocalin 2 as a Potential Ferroptosis-related Gene in Ulcerative Colitis. Inflamm Bowel Dis 29: 1446–1457, 2023.

46. Xin R, Zhang J, Zhang Y, Dai MP, Li Y, Ma JY, Shen M, Shen C, Chen Z, Zhu Q, et al: Pan-PPAR agonist bezafibrate alleviates psoriasis by suppressing LCN2-dependent ferroptosis. Free Radic Biol Med 247: 107–121, 2026.

47. Duggan E, Fuqua JD, Hagy B, Georgescu C, Miller BF, Van Remmen H and Brown JL: Phospholipid Glutathione Peroxidase Overexpression Mitigates Cancer Cachexia by Protecting Muscle Mass and Lowering Inflammation. J Cachexia Sarcopenia Muscle 17: e70255, 2026.

48. Zhang C, Wang H, Hu L, Zhang Q, Chen J, Shi L, Song X, Liu J, Xue K, Wang J, et al: Lipocalin-2 promotes neutrophilic inflammation in nasal polyps and its value as biomarker. Allergol Int 73: 115–125, 2024.

49. Choudhary BS, Chaudhary N, Khan BK, Vijan A, Mandal D, Pilankar L, Gawand S, Uttankar P, Sharma M, Shivashankar A, et al: LCN2 promotes focal adhesion formation and invasion by stimulating c-Src activation. J Cell Sci 138: 2025.

50. Jia K, Ding Y, Jin M, Yang Q, Wang X, Yan Y, Xiang J, Zhang R, Fang J, Jiang J, et al: Senescent endothelial cell-derived extracellular vesicles promote neoadjuvant chemoresistance in colorectal cancer via GPX4. Commun Biol 2026.

51. Yao F, Deng Y, Zhao Y, Mei Y, Zhang Y, Liu X, Martinez C, Su X, Rosato RR, Teng H, et al: A targetable LIFR-NF-kappaB-LCN2 axis controls liver tumorigenesis and vulnerability to ferroptosis. Nat Commun 12: 7333, 2021.

52. Hu J, Li Y, Lian B, Mao Y and Zhao L: Mechanism and role of regulated cell death in tumor immunity and immunotherapy. Cancer Commun (Lond) 45: 1456–1495, 2025.

53. Zhang T, Gu F, Lin W, Shao H, Jiang A and Guan X: Boosting cancer immunotherapy: drug delivery systems leveraging ferroptosis and immune checkpoint blockade. Front Immunol 16: 1611299, 2025.

