## Supplementary material for "STRIP2 promotes colorectal cancer malignant progression via stabilizing LCN2 to suppress ferroptosis": Western blot raw images

**Supplementary:** Uncropped Western blot raw images

The protein marker used is Tanon™ Prestained Protein Marker. (Cat# 180-6006. Shanghai Tanon Life Science Co.,Ltd.) in 10% gel. The typical protein marker pattern is as illustrated below for Western blot images.

To optimize antibody use, we cut the membranes to the target protein molecular weight range before antibody incubation. This approach effectively avoids unnecessary antibody consumption incurred by incubating the entire membrane. It is a common and recognized optimization in Western blot experiments when antibody resources are limited.

Chemiluminescent imaging was performed using an SCG-W2000 system (Wuhan Sevier Biotechnology Co., Ltd., Wuhan, China). However, it can only produce black-and-white images.

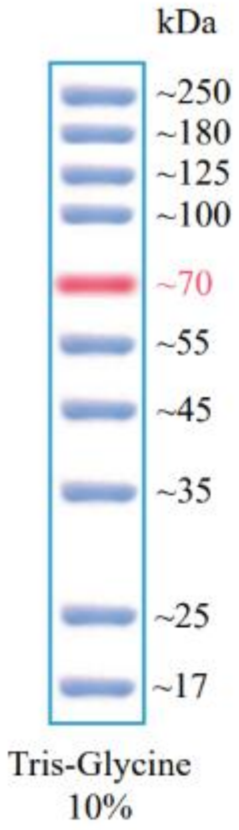


Tanon™ Prestained Protein
Marker for Western blot gels.

Figure 1D. Western blot analysis of STRIP2 protein levels in paired CRC tumor (T) and matched adjacent normal (N) tissues (n=5).

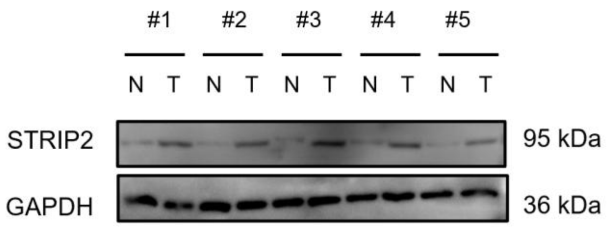


Raw Western blot images

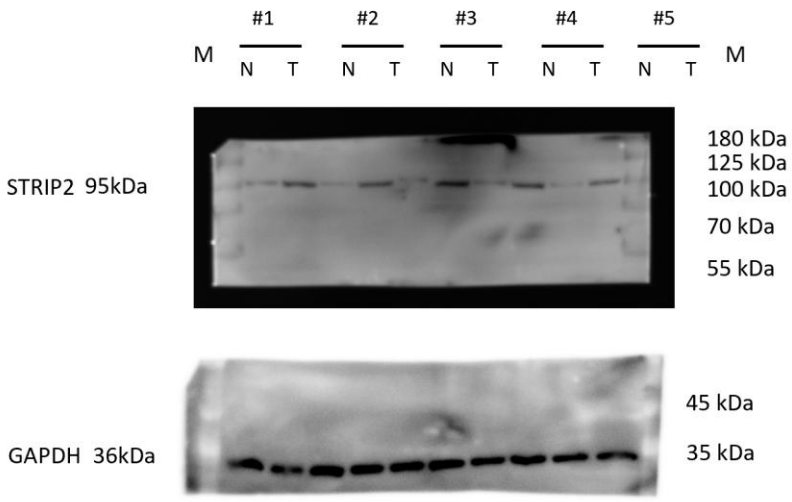

Figure 2B. Western blot analysis confirming STRIP2 protein silencing in SW480 and HCT116 cells (n=3).

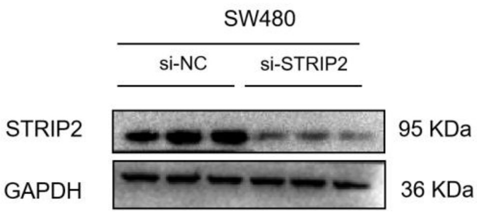


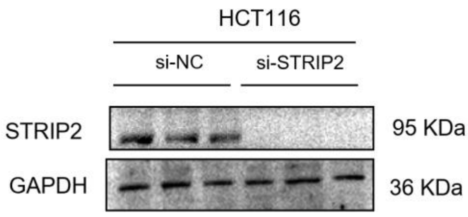


Raw Western blot images

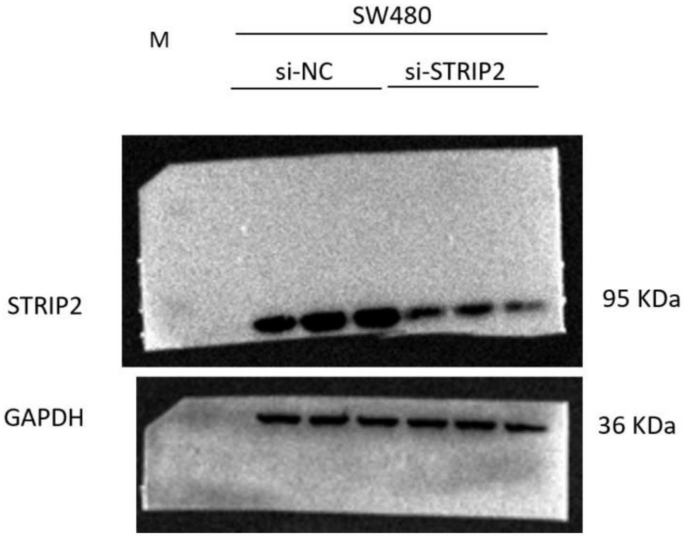


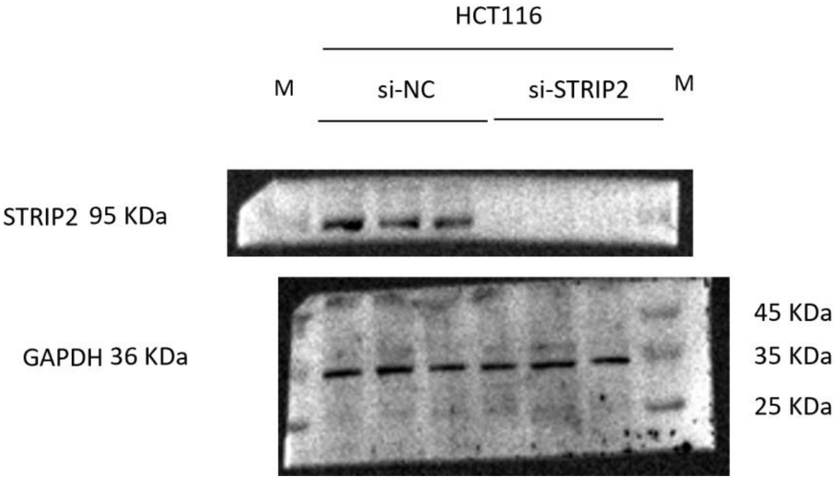

Figure 3E. Western blot analysis of epithelial–mesenchymal transition (EMT) marker proteins in STRIP2-knockdown SW480 and HCT116 cells (n=3).

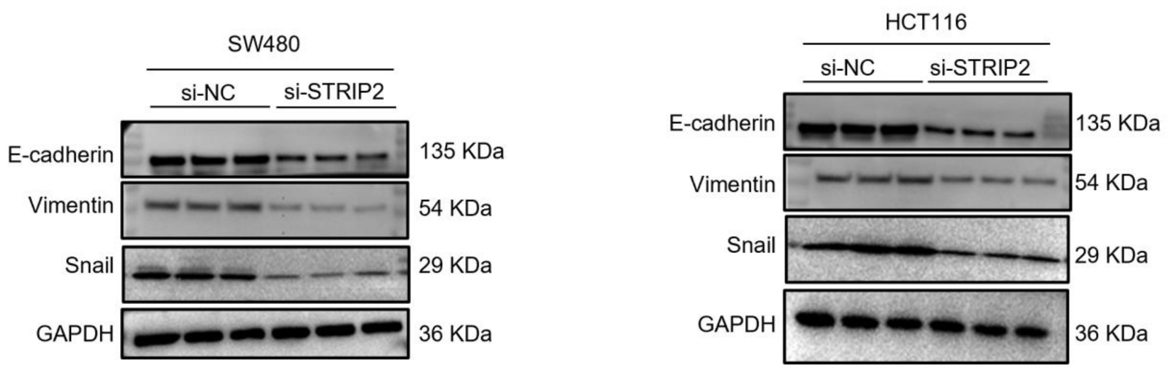


Raw Western blot images for SW480 cells

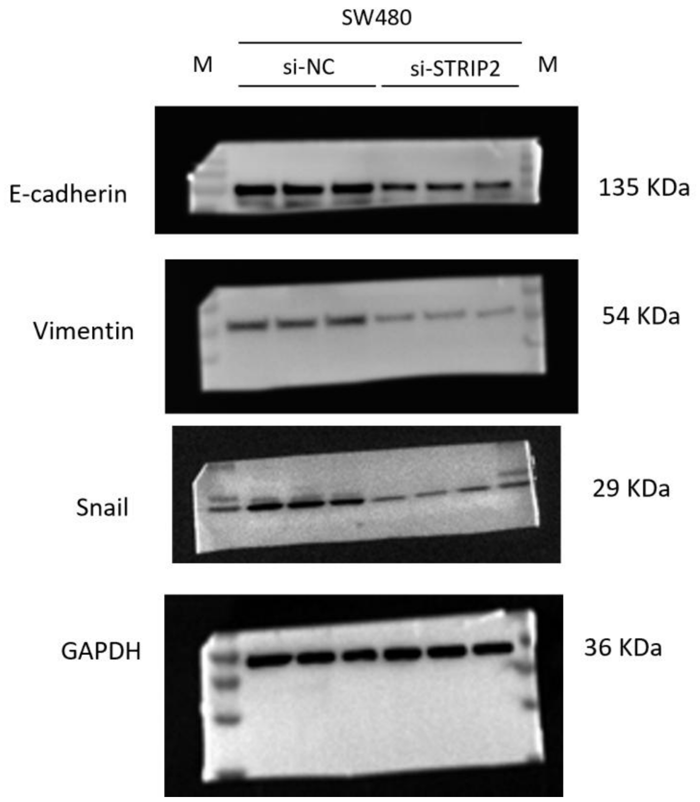


Raw Western blot images for HCT116 cells

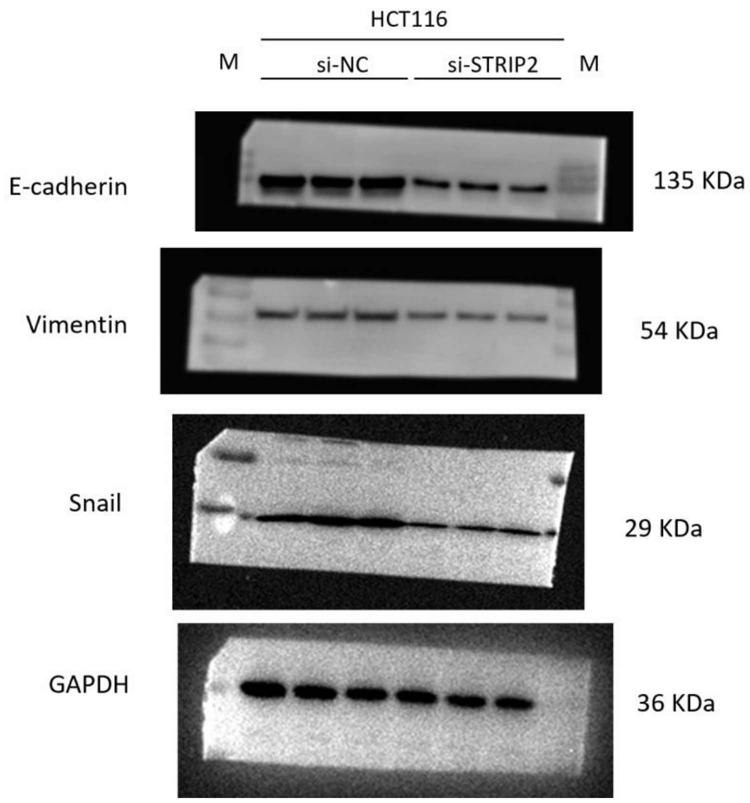

Figure 3G. Western blot analysis of apoptosis-related protein levels (n=3) in SW480.

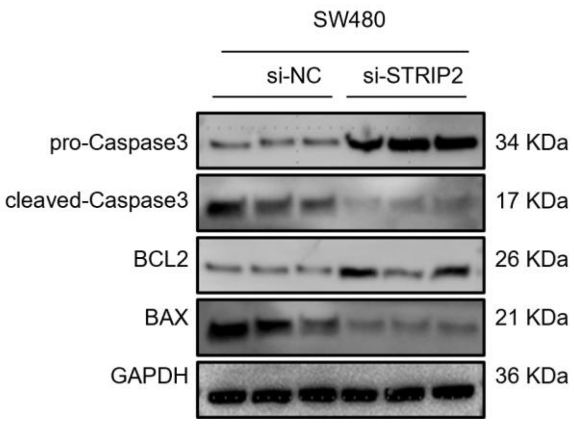


Raw Western blot images for SW480 cells

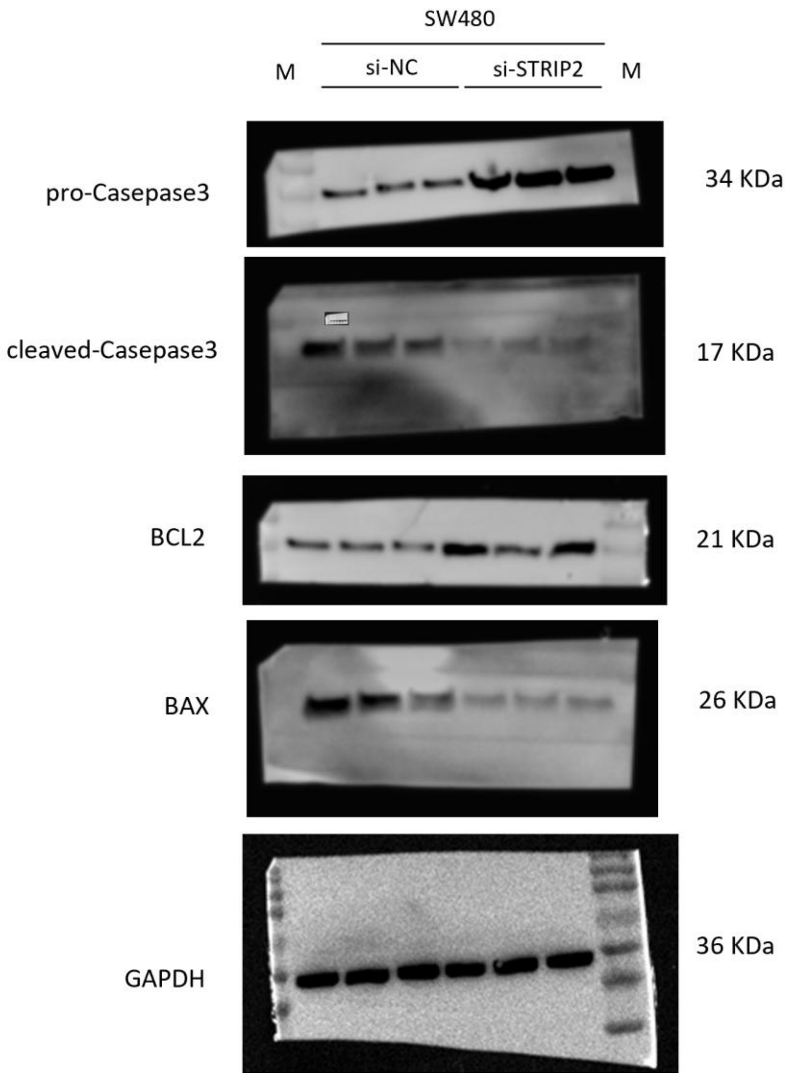

Figure 3G. Western blot analysis of apoptosis-related protein levels (n=3) in HCT116.

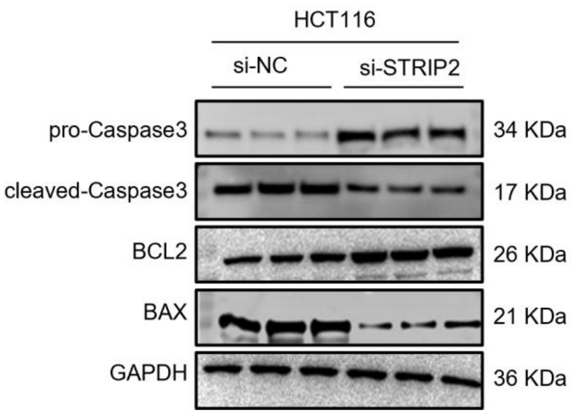


Raw Western blot images for HCT116 cells

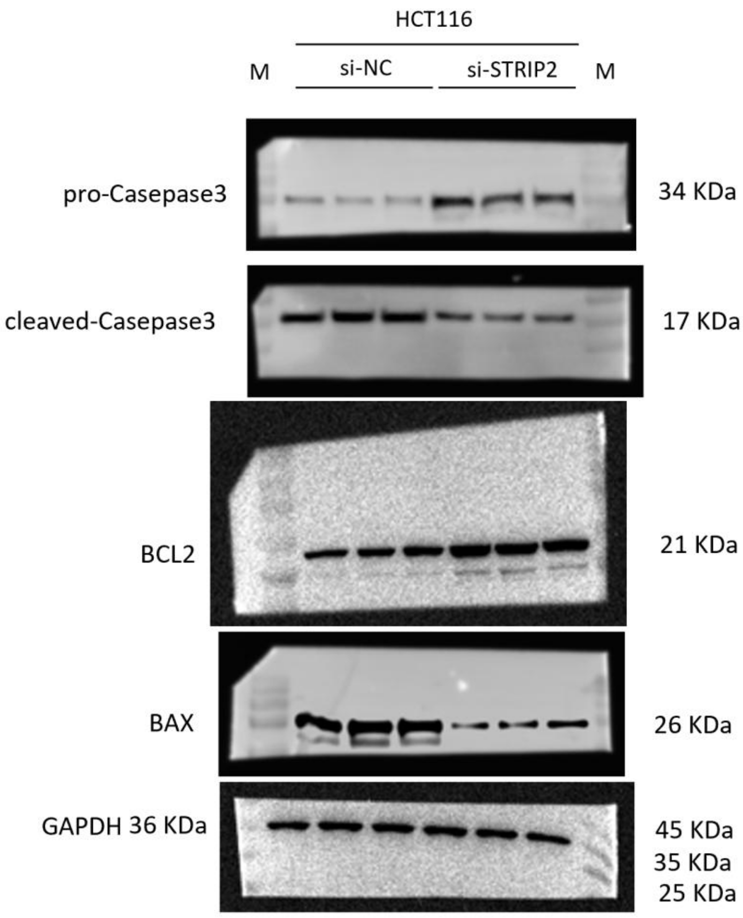

Figure 6A. Western blot of LCN2 in si-NC and si-STRIP2 SW480 and HCT116 cells.

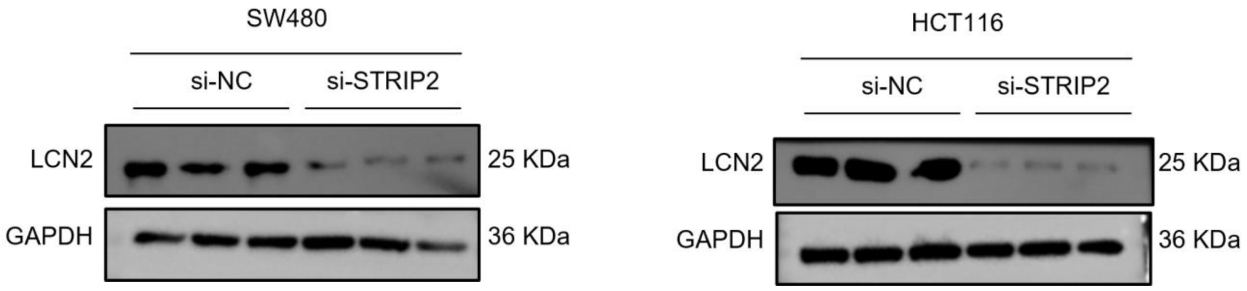


Raw Western blot images

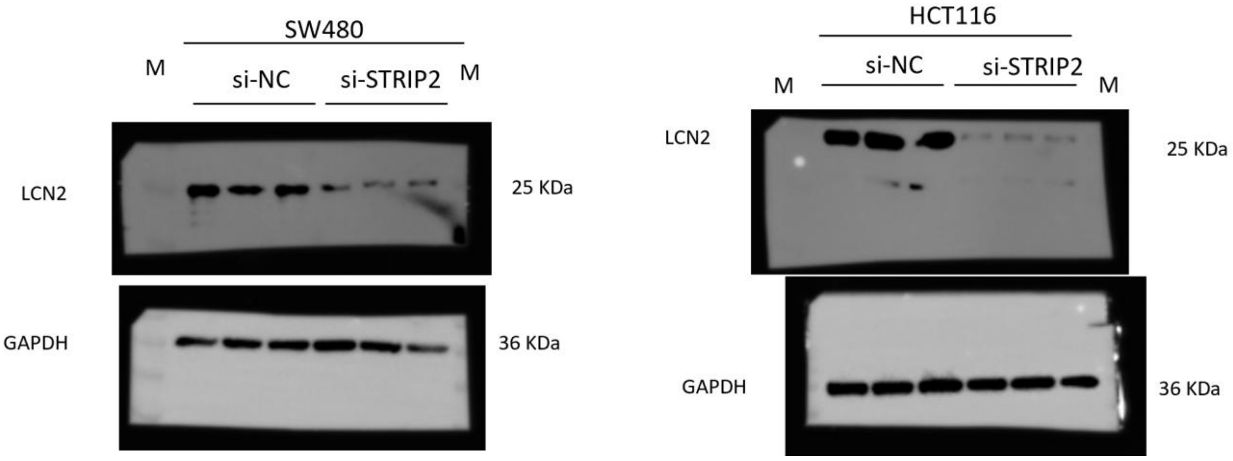


Figure 6B. Western blot of LCN2 and STRIP2 in 293T/17 cells with empty vector, single or combined LCN2/STRIP2 overexpression plasmids, with/without 6 h MG132 (20 μmol/l) treatment.

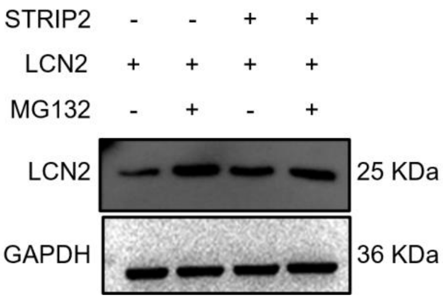


Raw Western blot images

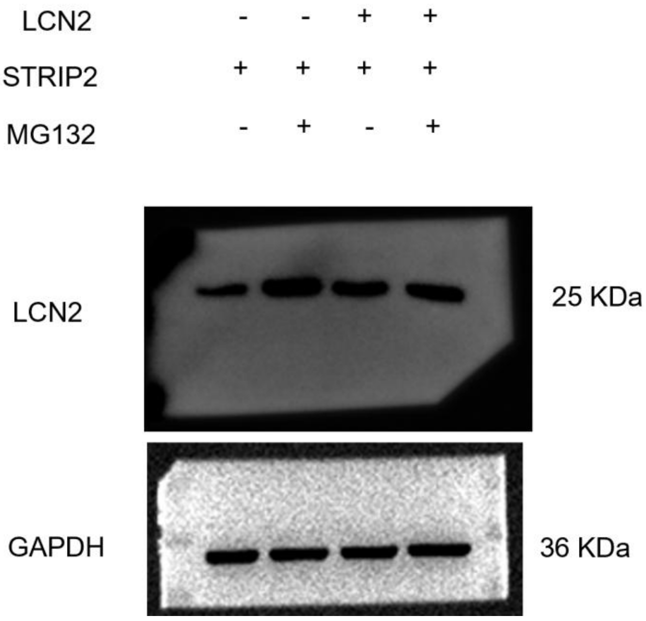

Figure 6C. Time-course western blot measuring LCN2 half-life in si-NC and si-STRIP2 HCT116 cells treated with CHX (100 μg/ml) for 0–15 h.

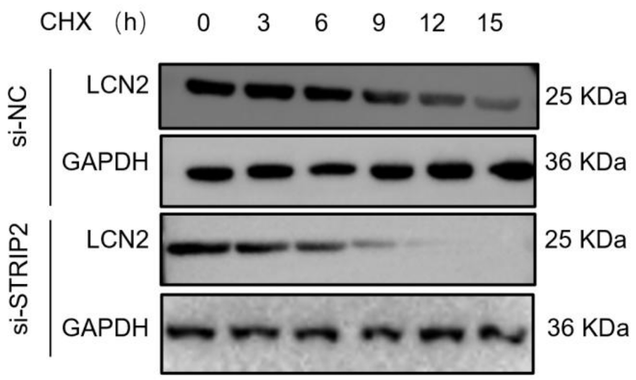


Raw Western blot images

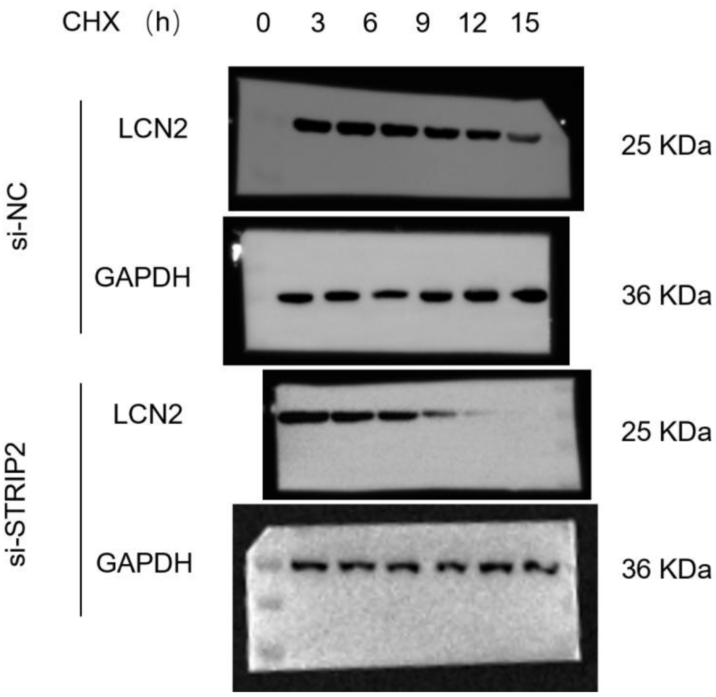

Figure 6D. Western blot detecting exogenous LCN2 and STRIP2 in co-transfected 293T/17 cells.

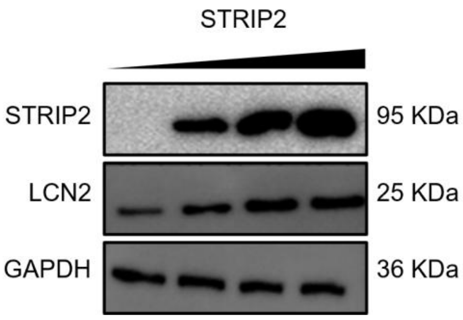


Raw Western blot images

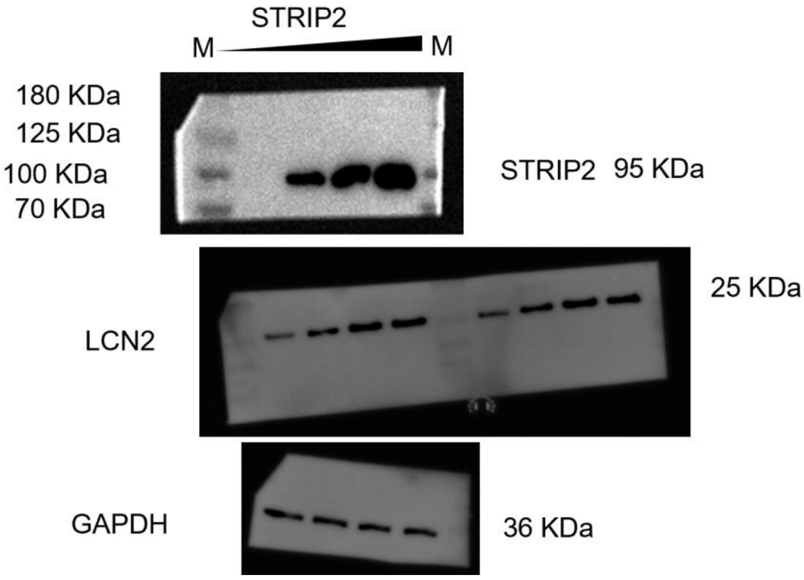

Figure 6E. Co-IP assay verifying STRIP2-mediated inhibition of LCN2 ubiquitination.

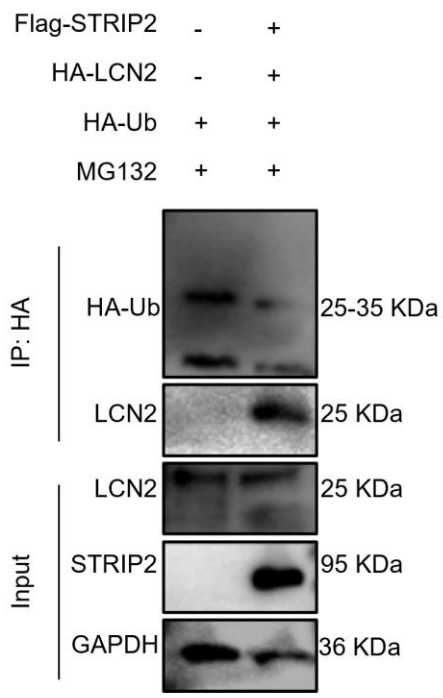


Raw Western blot images

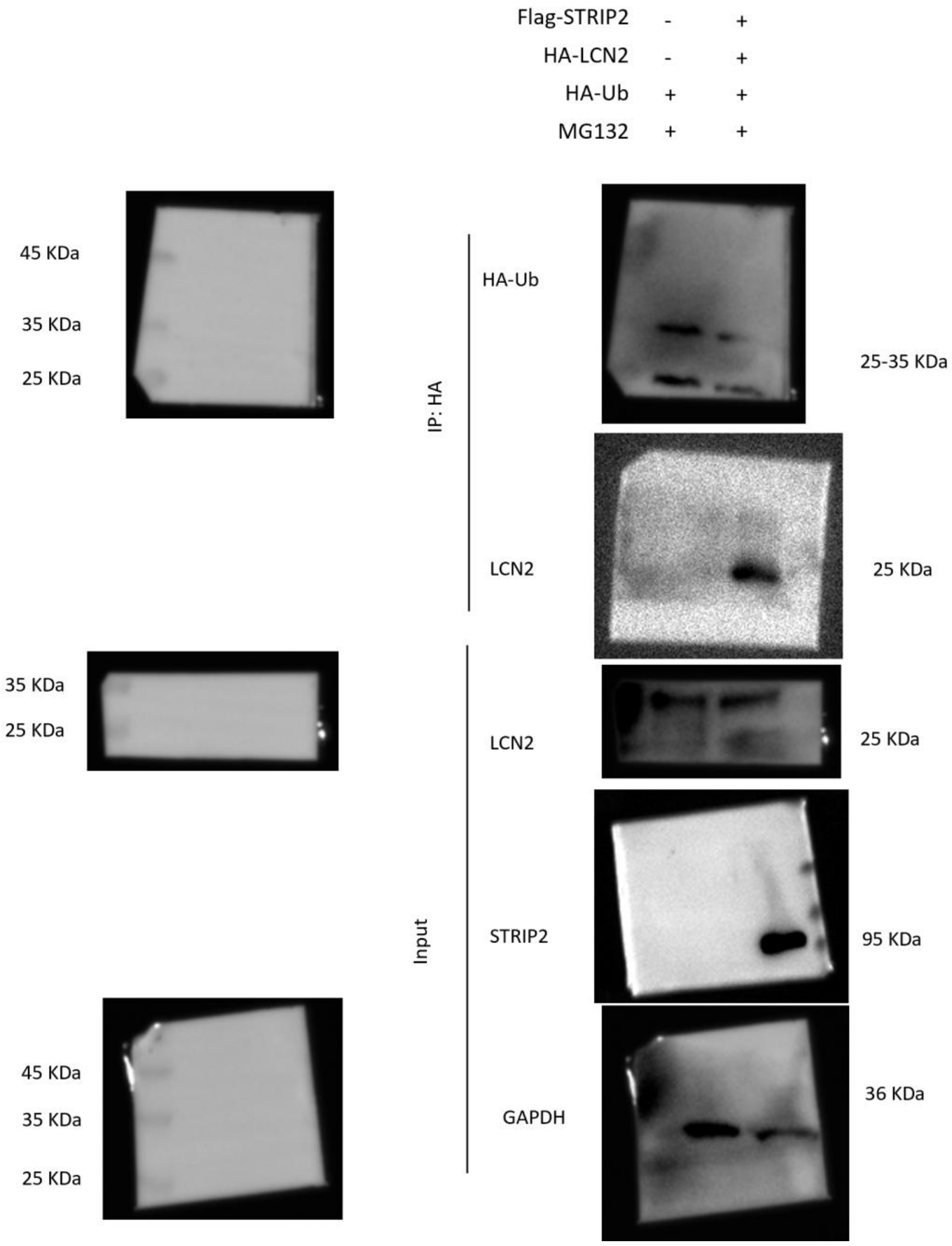

Figure 7A. Western blot of core ferroptosis regulatory proteins (ACSL4, SLC7A11, GPX4, FSP1 and NRF2) regulators in oe-NC, oe-STRIP2, si-NC and si-STRIP2 HCT116 cells (n = 3).

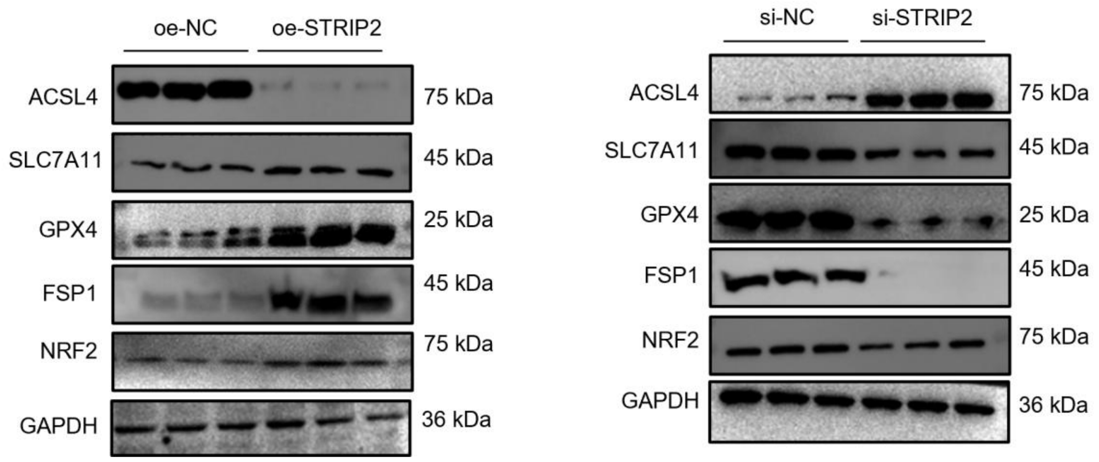


Raw Western blot images

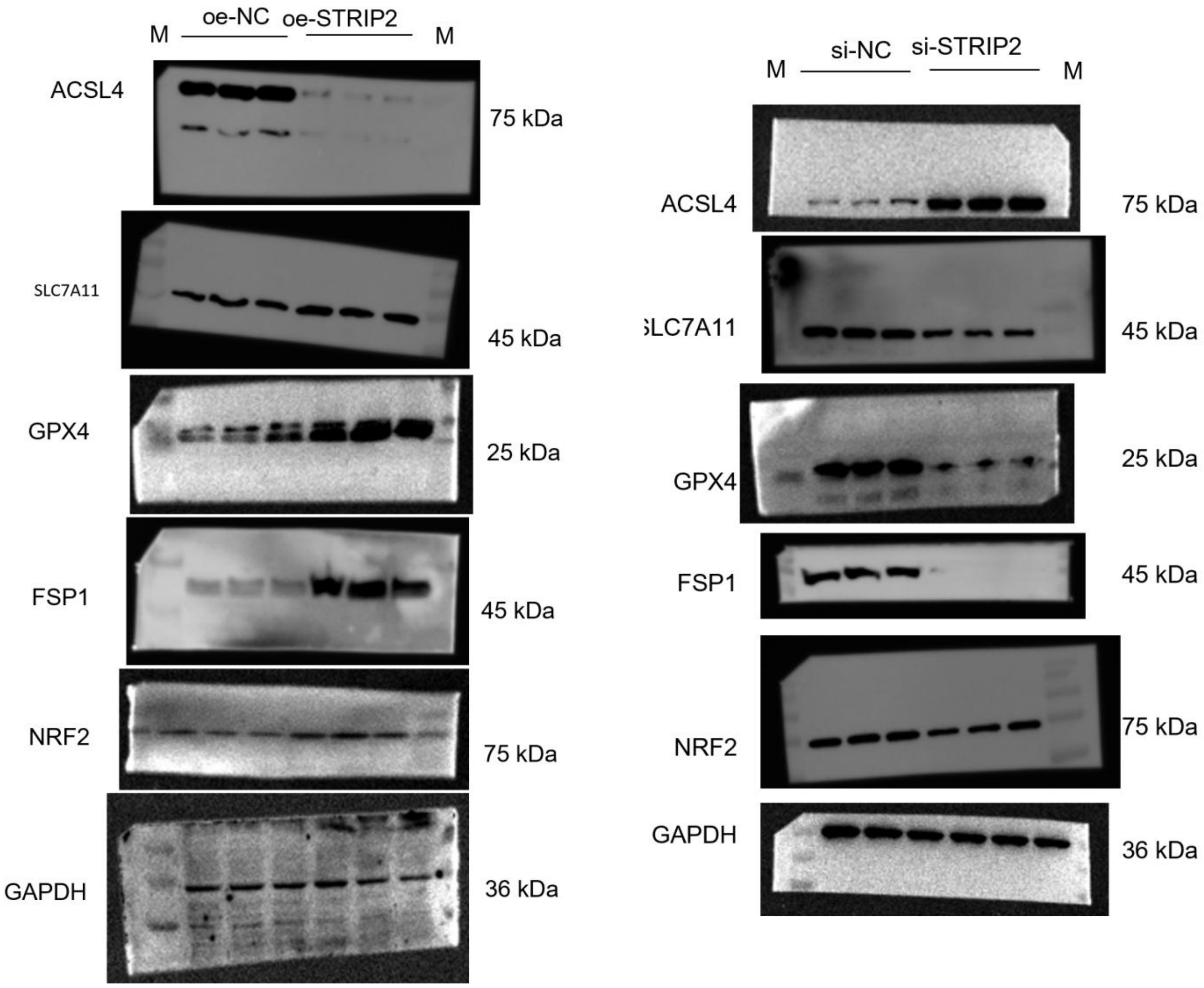
